# Comparative analysis of antibody- and lipid-based multiplexing methods for single-cell RNA-seq

**DOI:** 10.1101/2020.11.16.384222

**Authors:** Viacheslav Mylka, Jeroen Aerts, Irina Matetovici, Suresh Poovathingal, Niels Vandamme, Ruth Seurinck, Gert Hulselmans, Silvie Van Den Hoecke, Hans Wils, Joke Reumers, Jeroen Van Houdt, Stein Aerts, Yvan Saeys

**Affiliations:** VIB Tech Watch, VIB Headquarters, Ghent, Belgium; VIB Center for Brain & Disease Research, Leuven, Belgium; Department of Human Genetics, KU Leuven, Leuven, Belgium; Data mining and Modelling for Biomedicine, VIB Center for Inflammation Research, Ghent, Belgium; Department of Applied Mathematics, Computer Science and Statistics, Ghent University, Ghent, Belgium; Discovery Sciences, Janssen Research & Development, Pharmaceutical Companies of Johnson & Johnson, Beerse, Belgium

**Keywords:** hashing, scRNA-seq, MULTI-seq, CITE-seq, sample multiplexing

## Abstract

Multiplexing of samples in single-cell RNA-seq studies allows significant reduction of experimental costs, straightforward identification of doublets, increased cell throughput, and reduction of sample-specific batch effects. Recently published multiplexing techniques using oligo-conjugated antibodies or - lipids allow barcoding sample-specific cells, a process called ‘hashing’. Here, we compare the hashing performance of TotalSeq-A and -C antibodies, custom synthesized lipids and MULTI-seq lipid hashes in four cell lines, both for single-cell RNA-seq and single-nucleus RNA-seq. Hashing efficiency was evaluated using the intrinsic genetic variation of the cell lines. Benchmarking of different hashing strategies and computational pipelines indicates that correct demultiplexing can be achieved with both lipid- and antibody-hashed human cells and nuclei, with MULTISeqDemux as the preferred demultiplexing function and antibody-based hashing as the most efficient protocol on cells. Antibody hashing was further evaluated on clinical samples using PBMCs from healthy and SARS-CoV-2 infected patients, where we demonstrate a more affordable approach for large single-cell sequencing clinical studies, while simultaneously reducing batch effects.

## INTRODUCTION

Recent advances in single-cell and single-nucleus RNA sequencing (scRNA-seq and snRNA-seq) have had an unprecedented impact on our understanding of heterogenous cell populations (Aizarani et al. 2019; Davie et al. 2018; Han et al. 2018; Van et al. Hove 2019; Lake et al. 2016; Schaum et al. 2018). Current scRNA-seq experiments make it possible to routinely assay many thousands of cells at once, with recent datasets reporting hundreds of thousands to millions of cells (Davie et al. 2018; Park et al. 2020; Schaum et al. 2018). In standard single-cell workflows, individual samples need to be processed in parallel, which limits the throughput, increases reagent costs and has the potential to introduce batch effects. Recently, several approaches for multiplexing have been described, including the use of pre-existing genetic diversity (Kang et al. 2018) or by introducing sample-specific barcodes using oligo-labelled antibodies (Stoeckius et al. 2018), oligo-labelled lipid anchors (McGinnis et al. 2019), chemical labelling with oligos (Gehring; et al. 2018), or genetic cell labelling (Guo et al. 2019). Multiplexing samples by labelling cells or nuclei with sample-specific barcodes before pooling and single-cell compartmentalization, a technique called ‘hashing’, allows for accurate detection of two (doublets) or more (multiplets) cells originating from different samples but captured in the same compartment, which inevitably occurs in standard single-cell workflows. Therefore, implementing a barcoding multiplexing paradigm allows users to drastically increase the number of cells or nuclei loaded per reaction, which consequently decreases per-cell library preparation cost.

The development of oligo-labelled antibodies directed against cell surface proteins for sample multiplexing, is a direct evolution from the Abseq (Shahi et al. 2017), REAP-seq (Peterson et al. 2017) and CITE-seq (Stoeckius et al. 2017) protocols. One of the most widely used methods to date for detection of the cell epitome is by using the TotalSeq antibodies from Biolegend in combination with the scRNA-seq technologies from 10x Genomics. There are several types of TotalSeq antibodies to be used for cell labelling, including TotalSeq-A antibodies that contain a poly-A sequence mimicking a natural mRNA. These are designed to work with any sequencing platform that relies on poly-dT oligonucleotides as the mRNA capture method, while TotalSeq-B and TotalSeq-C antibodies contain a capture sequences that are compatible with the 10x Genomics 3’ scRNA-seq (v3 or v3.1) and 5’ scRNA-seq workflows, respectively. For human samples, the pre-mixed TotalSeq hashtag reagents recognize cell surface markers CD298 and β2-microglobulin. The success of using antibodies for hashing depends on the ubiquitous expression of these target antigens, which can be problematic for some samples or species (Federico Garrido 2019; Hermiston, Xu, and Weiss 2003), limiting the sample-agnostic, universal application of this method. An elegant way to overcome this limitation is the use of lipid anchors that are antigen independent and insert universally into the cell or nucleus membrane, irrespective of sample type (McGinnis et al. 2019). Both antibody-based and lipid-based methods are simple, straightforward and generally applicable to a wide range of single cell applications and platforms, while genetic cell labelling and chemical labelling with oligos tend to be more challenging. It is still unclear which method is most accurate in separating samples based on the inserted hashtags. In terms of labour intensity both hashing methods are comparable.

In this study, we compared antibody-based and lipid-based sample barcoding methods by multiplexing four distinct human cancer cell lines. By exploiting the intrinsic genetic variations of these cell lines, demultiplexing by genetic diversity serves as a ‘ground truth’ and allows determining the hashing accuracy of each method.

Single-cell suspensions can only be prepared from fresh tissues, which is a major roadblock for analysing clinical samples, archived materials and tissues such as the brain, for which cells cannot be readily extracted (Grindberg et al. 2013; Habib et al. 2016; Lake et al. 2016). To overcome these limitations, single-nuclei can be extracted and analysed similarly to a standard single-cell workflow (Habib et al. 2017). Therefore, we included nuclei samples in our comparison of hashing methods. Finally, we evaluated hashing accuracy of human PBMCs from different healthy and diseased patients using TotalSeq-A antibodies, which can be very relevant for single-cell sequencing in clinical studies.

## METHODS

### Cell Culture

MCF7, PC3, DU145 and MDA-MB-231 cells were maintained according to standard procedures in RPMI-1640 (Gibco, #21875034), F-12K (LGCstandards, ATCC 30-2004), EMEM (LGCstandards, ATCC 30-2003) and DMEM: F-12 (LGCstandards, ATCC 30-2006) medium respectively, supplemented with 10% fetal bovine serum (Gibco, #10082147) and 1% penicillin/streptomycin (100 U/ml and 100 μg/ml, respectively, Gibco, #15140122) at 37 °C with 5% CO2. RPMI-1640 medium used to culture the MCF7 cells was additionally supplemented with 0.01 mg/ml human recombinant insulin (Sigma Aldrich, #I3536).

### Single cell dissociation

To prepare single cell solutions of the cultured cell lines, the culture medium was removed, and the cells were washed with 1X PBS. Afterwards, the cells were trypsinized (0.05% trypsin-EDTA, Gibco, # 25300054) and pelleted at 200 × g for 5min. The cells were resuspended in 2 ml of culture medium, gently mixed with 8 ml of 1X PBS and put on ice. Afterwards, the cells were pelleted at 200 × g for 5min on 4°C and resuspended in 1 ml 1X PBS with 0.04% BSA. Cells were passed through a 40 μm cell strainer (Corning, # CLS431750-50EA) and counted with a LUNA FL counter, using 1 μl of acridine orange and propidium iodide dye (F23001, Westburg).

### Single nuclei dissociation

Single nuclei suspensions were prepared using a modified nuclei isolation protocol from 10x Genomics (demonstrated protocol: isolation of nuclei from single cell suspensions, #CG000124, revD). We replaced the original lysis buffer with Nuclei EZ Prep buffer (Sigma Aldrich, # NUC101-1KT) with a total lysis time of 5 min and proceeded according to the referenced protocol. Loss of cell viability as a proxy for nuclei extraction quality and the number of nuclei were assessed with a LUNA FL counter, using 1 μl of acridine orange and propidium iodide dye (F23001, Westburg).

### Antibody hashing

Cell and nuclei labelling were performed according to an adapted Biolegend cell hashing protocol (TotalSeq™-A Antibodies and Cell Hashing with 10x Single Cell 3’ Reagent Kit v3 3.1 Protocol, Biolegend). Briefly, cells were then incubated for 10 min with Fc receptor block (PN 422301, TruStain FcX, BioLegend) to block nonspecific antibody binding. Subsequently, cells were incubated with mixtures of barcoded antibodies for 30 min at 4 °C. In the TotalSeq-A experiment, MCF7, PC3, DU145 and MDA-MB-231 cells were labelled with 0.1 μg TotalSeq-A0251, -A0252, -A0253 and -A0254, respectively. For TotalSeq-C hashing, MCF7, PC3, DU145 and MDA-MB-231 cells were labelled with 0.1 μg TotalSeq-C0251, -C0252, -C0253 and -C0255, respectively. For both TotalSeq-A and TotalSeq-C experiments 200 000 cells of each cell line were labelled with the antibodies in 100 μl PBS containing 1% BSA. After labelling, cells were washed 3 times by resuspension in 1.4 ml PBS containing 1% BSA, followed by centrifugation (200g 5 min at 4 °C). After the final wash, cells were resuspended in PBS containing 0.04% BSA and equal amounts of labelled MCF7, PC3, DU145 and MDA-MB-231 cells were pooled before loading onto the 10x Chromium Single Cell B Chip (PN# 2000060) in case of TotalSeq-A hashing or onto the 10x Chromium chip A (PN# 2000019), because of the TotalSeq-C antibody compatibility with 5’ 10x chemistry (Chromium Single Cell Immune Profiling).

Nuclei were labelled with oligo-tagged anti-nucleoporin antibodies (#Mab414, Biolegend) (Sup. Table 1). MCF7, PC3, DU145 and MDA-MB-231 nuclei (840 000 of each line) were labelled with 0.5 μg TotalSeq-A0456, -A0457, -A0458 and -A0459, respectively, in 2% BSA solution with 0.02% Tween-20, 10mM Tris, 146mM NaCl, 1mM CaCl_2_, 21mM MgCl_2_ (Gaublomme et al. 2019). After the last wash nuclei were resuspended in the same labelling buffer as above with lower concentration of BSA (1%) and pooled prior to loading on chip B from 10x Genomics.

**Table 1.** Summary of CellRanger output.

| Hashing experiment | Estimated Number of Cells | Mean gene reads per cell | Hashtag Reads Usable* per Cell | Hashtag Median UMIs per Cell |
| --- | --- | --- | --- | --- |
| 1. TotalSeq-A cells | 11869 | 26182 | 1306 | 428 |
| 2. TotalSeq-A cells rep2 | 17611 | 14207 | 865 | 251 |
| 3. TotalSeq-C cells | 9229 | 31711 | 2934 | 1104 |
| 4. LMO (MULTI-seq) cells | 16827 | 26062 | 970 | 271 |
| 5. LMO (custom) cells | 21813 | 13764 | 1063 | 457 |
| 6. CMO nuclei | 15404 | 20396 | 536 | 426 |
| 7. TotalSeq-A nuclei | 23451 | 13345 | 267 | 207 |
| 8. TotalSeq-A nuclei rep2 | 9868 | 30371 | 520 | 258 |
| 9. TotalSeq-A PBMC1 (healthy) | 14635 | 23714 | 2195 | 712 |
| 10. TotalSeq-A PBMC2 (SARS-CoV-2) | 11372 | 27884 | 1289 | 584 |

\* reads that contain a recognized hashtag barcode, a valid UMI, and a cell-associated barcode.

The DNA library preparation was performed according to the manufacturer’s guidelines: CG000186 Rev A protocol for 5’ 10x Genomics chemistry and CG000185 Rev B protocol for 3’ 10x Genomics (v3 chemistry).

### Lipid hashing

Based on the MULTI-seq work, Lipid Modified Oligonucleotides (LMO) were used for cell hashing, and Cholesterol Modified Oligos (CMOs) were used for nuclei hashing. Labelling of nuclei was done with CMO because of the inefficacy of the LMOs to label nuclei in the presence of BSA. BSA specifically sequesters LMO and quenches LMO labelling (McGinnis et al. 2019).

In this work we have evaluated and compared the performance of commercially available oligos (custom LMOs purchased from Integrated DNA Technologies) with the LMOs used in the MULTI-seq work (McGinnis et al. 2019). Due to the commercial unavailability, LMOs with lower carbon chain length (C18; Stearyl modification) were used instead of the Lignoceric acid modified oligonucleotides (C22), described in the MULTI-seq manuscript.

The MULTI-seq lipid anchor and co-anchor were a kind gift from Christopher S. McGinnis at UCSF. The labelling of MCF7, PC3, DU145 and MDA-MB-231 cells was performed as previously described (McGinnis et al. 2019). Briefly, for each cell line 500k dissociated cells were washed and resuspended in PBS and incubated with 200 nM 1:1 molar ratio of the anchor and unique barcode oligonucleotides for 5 min on ice. Afterwards, each sample was incubated with 180nM of co-anchor in PBS on ice for 5min, followed by addition of 1% BSA in PBS to reduce off-target labelling when pooling. Cells were washed two times in 1% BSA in PBS after which all samples were pooled, filtered through a 40 μm cell strainer (Corning, # CLS431750-50EA), counted and loaded onto a single reaction of a 10x Chromium chip B.

The anchor and co-anchor of the commercial LMOs are available through Integrated DNA technologies. Also, the anchor and co-anchor of the CMOs were conjugated to cholesterol at the 3′ or 5′ ends via a triethylene glycol (TEG) linker and are also commercially available from Integrated DNA Technologies (**Sup. Table 2**). The labelling workflow for the commercially available LMOs and CMOs are identical to the MULTI-seq protocol (McGinnis et al. 2019).

**Table 2.** Hashing efficiency.

| Hashing experiment | Matching singlets |  |  |  |
| --- | --- | --- | --- | --- |
|  | MULTISeqDemux | SD | HTODemux | SD |
| 1. TotalSeq-A cells | **95.4%** | ± 0.4 | 94.2% | ± 1.6 |
| 2. TotalSeq-A cells rep2 | **90.9%** | ± 1.8 | 83.9% | ± 14.2 |
| 3. TotalSeq-C cells | **96.2%** | ± 0.9 | 93.9% | ± 2.6 |
| 4. LMO (MULTI-seq) cells | **84.9%** | ± 8.8 | 82.9% | ± 14.2 |
| 5. LMO (custom) cells | **68.5%** | ± 17.4 | 54.8% | ± 28.1 |
| 6. CMO nuclei | **84.1%** | ± 1.9 | 79.0% | ± 9.1 |
| 7. TotalSeq-A nuclei | 50.2% | ± 31.2 | **53.9%** | ± 19.9 |
| 8. TotalSeq-A nuclei rep2 | **63.3%** | ± 30.7 | NA<sup>#</sup> | NA<sup>#</sup> |
| 9. TotalSeq-A PBMC1 (healthy) | **84.1%** | ± 1.7 | 47.1% | ± 45.4 |
| 10. TotalSeq-A PBMC2 (SARS-CoV-2) | **83.6%** | ± 2.2 | 35.6% | ± 50.1 |

### Sequencing

The generated libraries were pooled targeting 85% mRNA and 15% hashtag oligo (HTO) and paired-end sequenced on individual lanes on a HiSeq4000 (Illumina) instrument, using the following read lengths: 28 bp Read1, 8 bp I7 Index and 91 bp Read2.

### PBMC sample preparation and single cell sequencing

PBMCs from 3 healthy control individuals and 3 SARS-CoV-2 patients were isolated as follows. Whole blood separation was performed by bringing whole blood, diluted with PBS 7.2 (ThermoFisher Scientific, # 20012027), in a Leucosep™ tube, (Greiner Bio-One, # 227290), prefilled with 15 mL Lymphoprep™(Stemcell technologies, # 07851), followed by a centrifugation step of 30 min at 400g (acceleration 5, brake 3). After isolation, the PBMCs were washed twice with PBS 7.2 and centrifuged at 350g during 10 min at 4°C. Isolated PBMCs were counted, cryopreserved in 1mL FCS/10%DMSO and stored in liquid nitrogen. After the thawing as described in the PBMC sample preparation protocol from 10x Genomics (CG00039, Rev C) cells were labelled according to the adapted CITE-seq protocol (Stoeckius et al. 2017). In brief, 1.25 × 10^6 PBMCs were spun down and resuspended in 25 μl CITEseq antibody cocktail (TotalSeq™-A Antibodies, BioLegend) containing among others one hashing antibody per each sample. After 30 min incubation on ice, cells were washed 3 times, pooled together (3 samples) and resuspended in PBS+0.04% BSA at a final concentration of 1000 cells/μl. Cells were counted using automated counter LUNA FL™ (Logos Biosystems). Cellular suspensions (target recovery of 10000 cells) were loaded on a 10x Genomics NextGEM Single-Cell Instrument, chip G (# 2000177) to generate single-cell Gel Bead-in-EMulsion (GEMs). Single-cell RNA-Seq libraries were prepared using GemCode Single-Cell V3.1 (NextGEM) 3′Gel Bead and Library Kit (10x Genomics) according to the manufacturer’s instructions. Sequencing libraries were sequenced with Illumina NovaSEQ S4 flow cell with custom sequencing metrics (single-indexed sequencing run, 28/8/0/98 cycles for R1/i7/i5/R2).

### Single-cell ata processing

#### Mapping

The raw sequencing data were demultiplexed and further converted into a single-cell level gene counts matrix with Cell Ranger 3.1.0 with default parameters (https://github.com/10xGenomics/cellranger). Only the number of expected cells was adapted in accordance with the number of targeted (captured) cells in each experiment. The mRNA reads were mapped to the human reference genome (assembly hg38 build 95) with the reads allowed to map to both exonic and intronic regions. For every type of hashing strategy, a specific Feature Reference File was generated and used as input for Cell Ranger. Filtered count matrices were used for downstream analysis.

### Freemuxlet-based demultiplexing

The deconvolution of the four cell lines was determined by a genotype-free tool freemuxlet which is an extension of demuxlet (Kang et al. 2018). Both tools are from the popscle (commit 7b141e3) software available at https://github.com/statgen/popscle/. To reduce computation time, the input bam file was sorted to contain only reads that i) overlap with the SNPs in the VCF file and ii) have a cell barcode listed in the cell barcode list. Code is available at https://github.com/aertslab/popscle_helper_tools. The filtered bam file was further used with the popscle dsc-pileup tool to pileup the reads and corresponding base quality for each overlapping SNPs and each barcode. The reference vcf file was assembled from the 1000 Genomes Project GRCh38 genetic variants data (Auton et al. 2015) adapted to the index used for mapping and the variant allele frequency > 0.9 and <0.1. were discarded. Next, freemuxlet with default parameters was used to determine sample identity and identify doublets.

### Sample demultiplexing and clustering

To assign hashing tags for each cell two strategies implemented in Seurat 3.1.4 package (Butler et al. 2018; Stuart et al. 2019) in R version 3.6.0 were evaluated on the filtered feature-barcode matrix generated by CellRanger: i) HTODemux function with default parameters and ii) incorporated into Seurat MULTIseqDemux function (McGinnis et al. 2019) with argument autoThresh = TRUE. Subsequent clustering and visualization of the data sets was performed in R using Seurat version 3.1.4 functions as described in the “Demultiplexing with hashtag oligos (HTOs)” vignette available at (https://satijalab.org/seurat/v3.1/hashing_vignette.html). The hashing efficiency was calculated as the overlap between cells annotated using Seurat (MULTISeqDemux and HTODemux functions) and freemuxlet (singlets only).

### Doublet identification

For each dataset, doublet cell detection was performed using DoubletFinder (v2.0.1) (Mcginnis et al. 2019) with the 10% as expected doublet rate and the following parameters: PCs=1:25, pN=0.25, pK= the highest value on BCmvn plot (find.pK function); Scrublet (v 0.1.3) and freemuxlet with default parameters, and Seurat (MULTISeqDemux (autoTresh=T) and HTODemux (default parameters) functions). The functions incorporated in Seurat predict doublets based on hashtag signal, while other tools use transcriptome data for this.

## RESULTS

### Cell hashing using antibodies and lipids accurately demultiplexes majority of cells

We first compared antibody-based and lipid-based hashing techniques by pooling four hashed cancer cell lines (MCF7, PC3, DU145 and MDA-MB-231) into a single 10x Chromium run followed by a single analysis pipeline (**Figure 1**) for each type of hashing. The hashing efficiency was calculated as an overlap between cell line annotation using Seurat (MULTISeqDemux and HTODemux functions) and freemuxlet as a reference annotation (**Table 2**), as follows: hashing efficiency (%) equals the number of all matching singlets (e.g. a given cell *i* annotated as one cell line by both Seurat and freemuxlet) divided by the number of all freemuxlet-annotated singlets.

**Figure 1.**
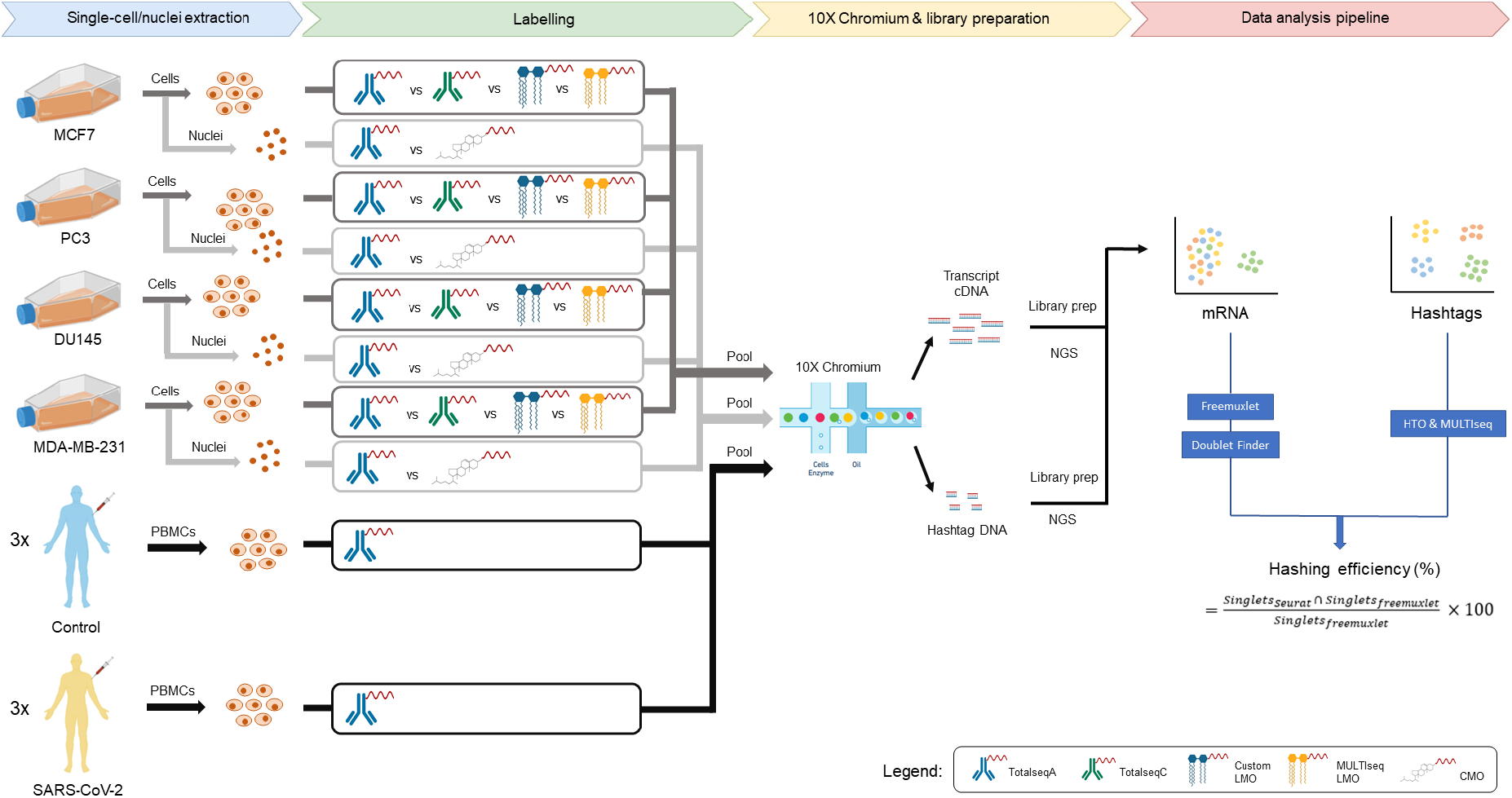
Overview of the experimental setup. Four cancer cell lines were used to extract cells or nuclei to process further with the different labelling methods. After pooling, the samples were run on a 10x Genomics Chromium platform and libraries were sequenced. Hashing efficiency was determined by demultiplexing the clusters based on single nucleotide polymorphisms (SNPs) and comparing those clusters to the respective hashtag clusters. LMO: lipid-modified oligonucleotides; CMO: cholesterol-modified oligonucleotides.

Importantly, the tested demultiplexing functions from Seurat use antibody- or lipid-derived hashtag oligo (HTO) expression data for cell line annotation, while freemuxlet uses transcriptome data and a Bayes Factors approach to evaluate the likelihood of a hashed droplet being a doublet and estimation of its cell line origin. Freemuxlet exploits a similar algorithm as demuxlet (Kang et al. 2018), but does not require externally genotyped data (popscle package).

One of the first characteristics evaluated for all types of hashing techniques in our study was the level of expression of each HTO. When comparing the expression of each antibody- or lipid-derived hashtag oligo overlaid on the Uniform Manifold Approximation and Projection (UMAP) derived from the gene expression data, we observed a distinct barcode for each of the four cell line clusters. Cells with more than one HTO generally group together in between the large clusters, suggesting that these groups of cells are indeed doublets/multiplets. (**Figure 2A**). Each cell cluster expresses an intrinsic gene marker relevant to each cell line (**Sup. Fig. 1**). These findings are corroborated when each cluster is identified based on inherent genetic variation using freemuxlet (**Figure 2B**), clearly separating the four cell lines and doublets. We also evaluated sample multiplexing on the same cells using 5’ scRNA-seq chemistry and compatible TotalSeq-C hashing antibodies (**Figure 3**). Cells demultiplexed using MULTISeqDemux had a high concordance (95 % for TotalSeq-A and 96% for TotalSeq-C hashing) with freemuxlet-based annotation (singlets only) (**Figure 2B, 3B** and **Table 2**). The custom lipid (LMO) and MULTI-Seq lipid hashing techniques demonstrated 68.5 % and 84.9 % hashing efficiency, respectively (**Table 2**). The MULTISeqDemux function annotated 1.1%, 1.8%, 21% and 7% of cells in TotalSeq-A, TotalSeq-C, custom lipid (LMO) and MULTI-Seq lipid experiments respectively, as “Negatives” (**Figure 2C, 3C, 4C, 5C**), which have a background expression for each hashtag according to the algorithm of the MULTISeqDemux function (Stoeckius et al. 2018)(**Figure 2D, 3D**). We observed ~2 times less unique molecular identifiers (UMIs) and ~3 times less genes per cell in “Negatives” compared to singlets in case of antibody hashing (**Figure 2D, 3D**). For the lipid hashing (LMOs) experiments this ratio was only 1.1 times less for both number of UMIs and genes per cell (**Figure 4D, 5D**). Even though the hashtag fractions of at least “TotalSeq-A cell” and “custom LMO cell” samples were sequenced to the same depth – around 400 hashtag reads per cell (**Table 1**). It’s noticeable that lower hashing efficiency when using lipids compared to TotalSeq antibodies coincides with higher number of cells in the lipid hashing samples (**Table 1, Sup. Fig. 2**). Nevertheless, the repeated TotalSeq-A experiment with an increased number of cells (17611) still demonstrated better hashing efficiency (90.9 %) even when compared to the MULTI-Seq lipid hashing (84.9 %, 16827 cells) (**Table 1** and **2**). Remarkably, the number of “Negatives” was relatively higher in DU145 cells hashed with both types of lipid hashes (custom and MULTI-seq) (**Figure 4C, 5C**), which is in line with the relatively low hashtag expression in this cell line (**Figure 4A, 5A**). This observation might point to a potentially lower affinity of the tested lipids with DU145 cells. When we compared doublet annotation by freemuxlet and MULTISeqDemux, we noticed that 2.4%, 6.3%, 37.9% and 19.1% of freemuxlet-annotated doublets in TotalSeq-A, TotalSeq-C, custom lipid (LMO) and MULTI-Seq lipid experiments respectively, were recognized as singlets by the MULTISeqDemux function (**Sup. Table 3**). We observed some variation across different hashing techniques in doublet annotation by different doublet detection tools including also scrublet and doubletFinder (**Sup. Figure 3, 4, 5**).

**Figure 2.**
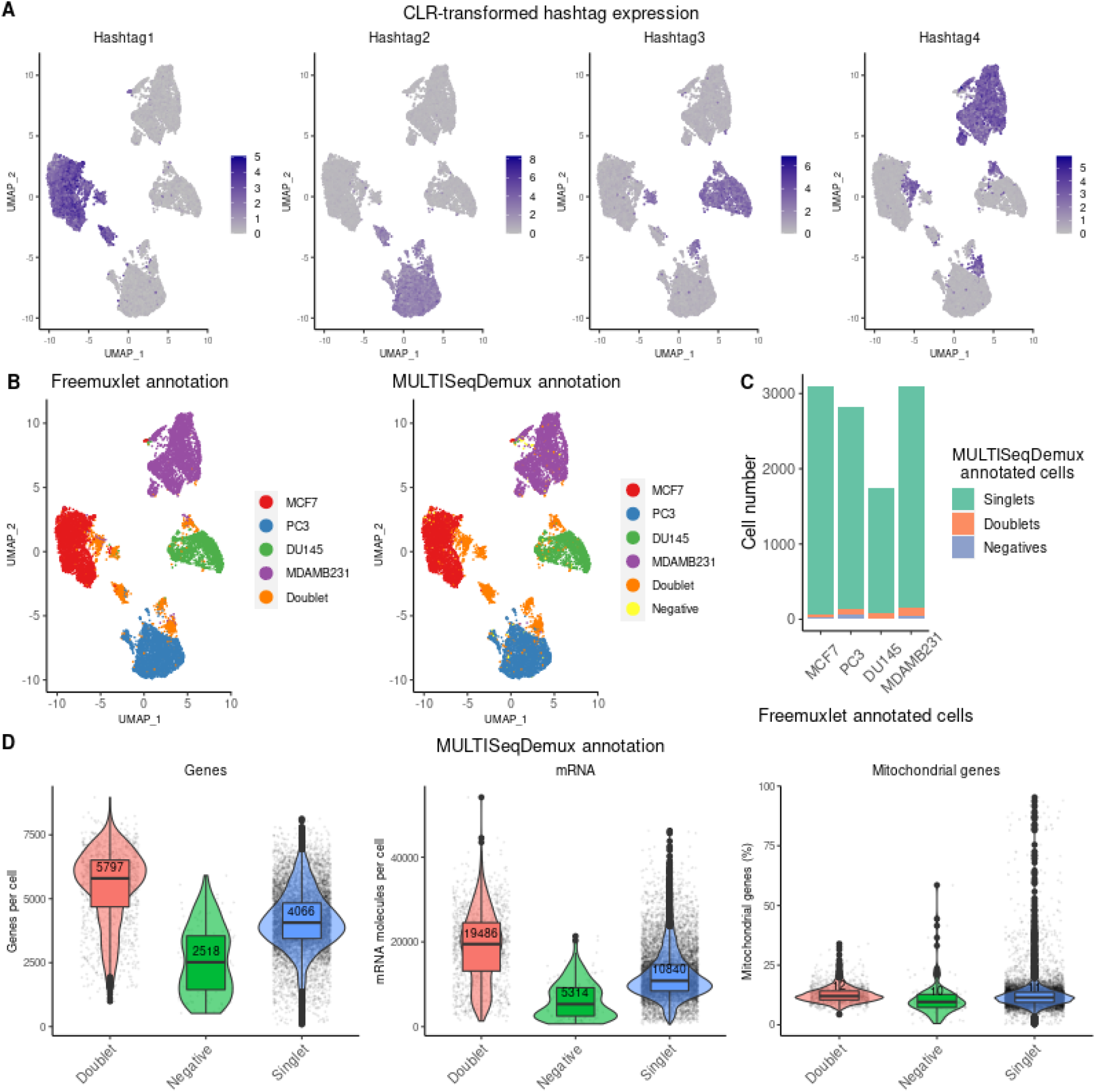
TotalSeq-A hashing on cells. **A.** Gene-cell matrices were generated using CellRanger, followed by log-transformation of gene UMI counts and cell clustering (gene expression, Principal Component Analysis (PCA) reduction) using Seurat. The hashtag UMI counts were CLR-transformed (centered log-ratio) and visualised in blue color on the gene expression UMAP plots. **B.** Cell annotation (4 cell lines) was performed using freemuxlet (gene expression) or Seurat (MULTISeqDemux function applied on hashtag counts data) and visualized on the gene expression UMAP plots. **C.** MULTISeqDemux-annotated singlets (MCF7, PC3, DU145 or MDAMB231) and negatives (cells with background expression for each hashtag) were matched with the freemuxlet-annotated cells (MCF7, PC3, DU145 or MDAMB231). The rest (unmatched) of freemuxlet-annotated singlets were assigned as doublets and altogether visualized in barplots. **D.** Gene expression were log-transformed using Seurat and detected genes (left plot), UMIs (middle plot) and percentage of mitochondrial genes expression (right plot) in cells were visualised as violin-box plots with median values highlighted in red, across MULTISeqDemux-annotated groups (singlets, doublets, negatives on basis of hashtag expression).

**Figure 3.**
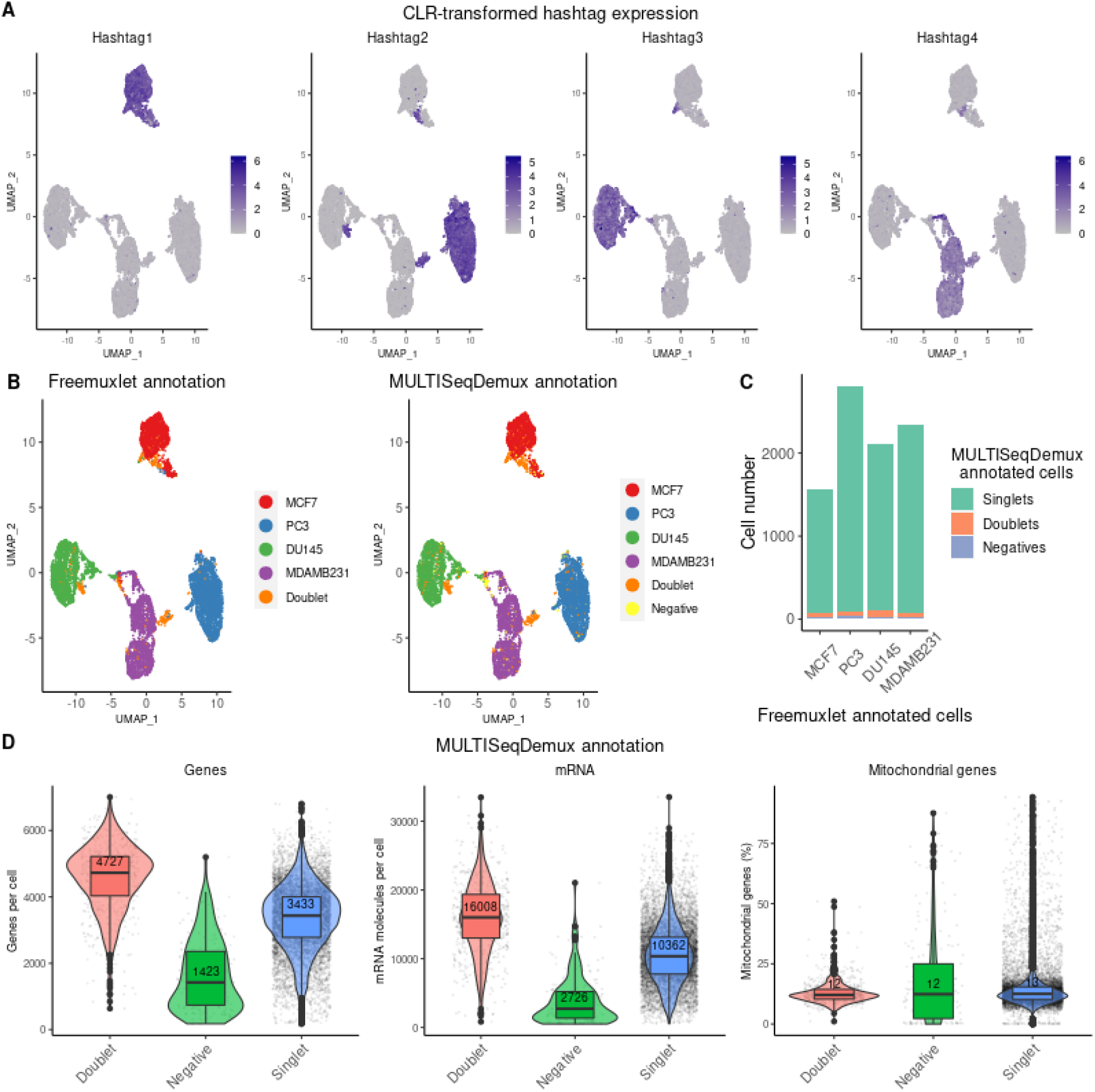
TotalSeq-C hashing on cells. **A.** Gene-cell matrices were generated using CellRanger, followed by log-transformation of gene UMI counts and cell clustering (gene expression, PCA reduction) using Seurat. The hashtag UMI counts were CLR-transformed and visualised in blue color on the gene expression UMAP plots. **B.** Cell annotation (4 cell lines) was performed using freemuxlet (gene expression) or Seurat (MULTISeqDemux function applied on hashtag counts data) and visualized on the gene expression UMAP plots. **C.** MULTISeqDemux-annotated singlets (MCF7, PC3, DU145 or MDAMB231) and negatives (cells with background expression for each hashtag) were matched with the freemuxlet-annotated cells (MCF7, PC3, DU145 or MDAMB231). The rest (unmatched) of freemuxlet-annotated singlets were assigned as doublets and altogether visualized in barplots. **D.** Gene expression were log-transformed using Seurat and detected genes (left plot), UMIs (middle plot) and percentage of mitochondrial genes expression (right plot) in cells were visualised as violin-box plots with median values highlighted in red, across MULTISeqDemux-annotated groups (singlets, doublets, negatives on basis of hashtag expression).

**Figure 4.**
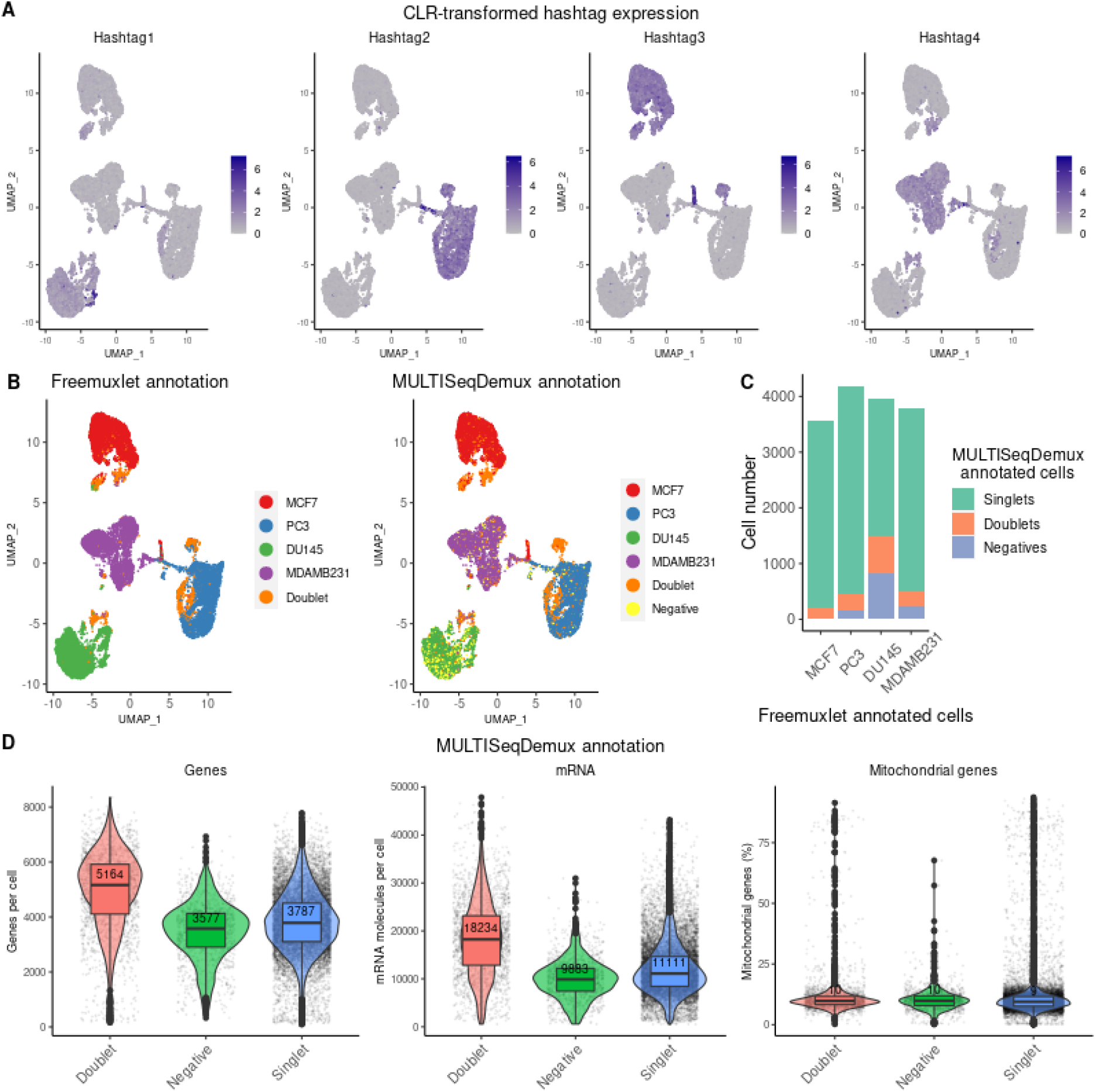
Lipid-modified oligo (LMO MULTI-seq) hashing on cells. **A.** Gene-cell matrices were generated using CellRanger, followed by log-transformation of gene UMI counts and cell clustering (gene expression, PCA reduction) using Seurat. The hashtag UMI counts were CLR-transformed and visualised in blue color on the gene expression UMAP plots. **B.** Cell annotation (4 cell lines) was performed using freemuxlet (gene expression) or Seurat (MULTISeqDemux function applied on hashtag counts data) and visualized on the gene expression UMAP plots. **C.** MULTISeqDemux-annotated singlets (MCF7, PC3, DU145 or MDAMB231) and negatives (cells with background expression for each hashtag) were matched with the freemuxlet-annotated cells (MCF7, PC3, DU145 or MDAMB231). The rest (unmatched) of freemuxlet-annotated singlets were assigned as doublets and altogether visualized in barplots. **D.** Gene expression were log-transformed using Seurat and detected genes (left plot), UMIs (middle plot) and percentage of mitochondrial genes expression (right plot) in cells were visualised as violin-box plots with median values highlighted in red, across MULTISeqDemux-annotated groups (singlets, doublets, negatives on basis of hashtag expression).

**Figure 5.**
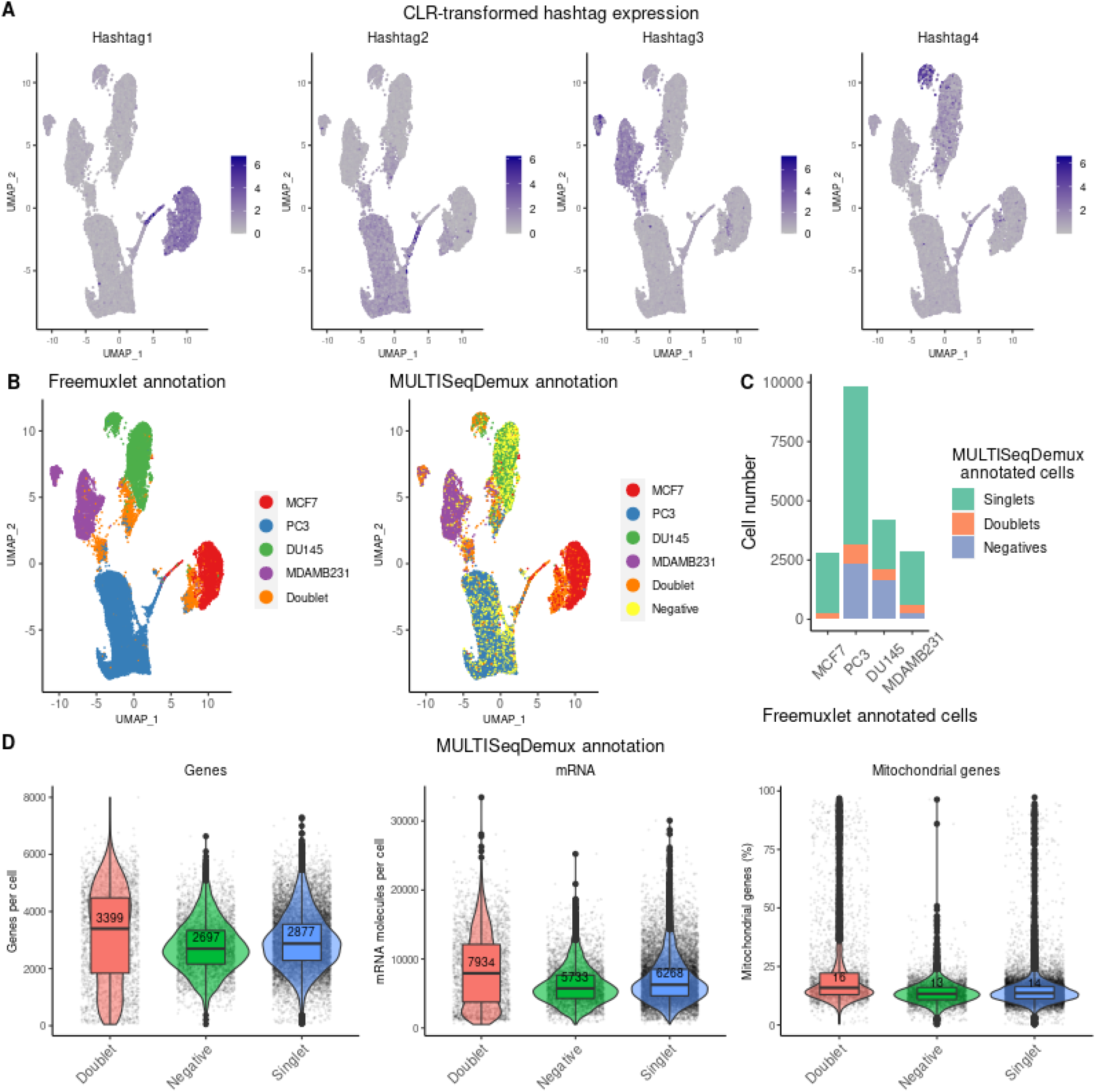
Lipid-modified oligo (custom LMO) hashing on cells. **A.** Gene-cell matrices were generated using CellRanger v3, followed by log-transformation of gene expression UMI counts and cell clustering using Seurat. The hashtag UMI counts were CLR-transformed and visualised in blue on the gene expression UMAP plots. **B.** Cell annotation (4 cell lines) was performed using freemuxlet (gene expression) or Seurat (MULTISeqDemux function applied on hashtag counts data) and visualized on the gene expression UMAP plots. **C.** MULTISeqDemux-annotated singlets (MCF7, PC3, DU145 or MDAMB231) and negatives (cells with background expression for each hashtag) were matched with the freemuxlet-annotated cells (MCF7, PC3, DU145 or MDAMB231). The rest (unmatched) of freemuxlet-annotated singlets were assigned as doublets and altogether visualized in bar plots. **D.** Gene expression data were log-transformed using Seurat and detected genes (left plot), UMIs (middle plot) and percentage of mitochondrial genes expression (right plot) in cells were visualised as violin-box plots with median values highlighted in red, across MULTISeqDemux-annotated groups (singlets, doublets, negatives on basis of hashtag expression).

When comparing the correct assignment of cells by the MULTISeqDemux function with their attributed genotypes, the percentage of hashtag swapping (fraction of each hashtag found among singlets in non-labelled cell lines) varied considerably between technologies with the lowest average value of ~ 0,1% mislabelled cells in both TotalSeq-A and TotalSeq-C hashing experiments, 0.89% mislabelled cells in the MULTI-Seq LMO hashing and 2.68% in the custom LMO hashing experiment (**Sup. Table 4**).

Regarding the performance of the demultiplexing functions from Seurat, we noticed that MULTISeqDemux function (autoTresh =T) overall correctly deconvolutes more cells than HTODemux function (default parameters) (**Table 2**).

Additional to the comparison of different hashing techniques on cells, we also estimated the level of expression of CD298 and β2-microglobulin in different human cells (mainly cancer cell lines) using flow cytometry. Both antigens are targeted by human TotalSeq -A and -C hashing antibodies and for their detection we used flow cytometry antibodies with the same clones. All the tested 17 human cell types including PBMCs, HEK293A and THP-1 cells express both antigens, potentially enabling antibody-based hashing on these cells (**Sup. Fig. 7**). To conclude, the Totalseq -A and -C hashing antibodies provide excellent hashing capabilities for a wide range of human sample types.

### Lipid (CMO)-based hashing outperforms antibody hashing on nuclei

For each of the four cell lines we extracted the nuclei to compare the antibody-based and lipid-based hashing efficiency. Nuclei hashing using lipids (cholesterol-modified oligos or CMO) and TotalSeq-A antibodies demonstrated overall lower signal-to-noise ratio of hashtag expression compared to TotalSeq antibody cell hashing (**Figure 6A, 7A** vs **Figure 2A, 3A**). Nevertheless, the nuclei hashed with cholesterol oligos followed by demultiplexing using MULTISeqDemux demonstrated 84% concordance with the reference annotation by freemuxlet (singlets correctly assigned to one of each 4 cell lines), or 79% when using HTODemux function (**Table 2**). TotalSeq-A-based nuclei hashing was less accurate (50% of all freemuxlet-annotated singlets for MULTISeqDemux and 54% for HTODemux function), mainly due to a high number of nuclei recognized as “Negatives” (cells with a background signal for each hashtag) by the demultiplexing functions used for this benchmarking study (**Figure 7B, 7C**). It was especially prominent in the MCF7 cell cluster (**Figure 7B, 7C**). However, it was possible to improve the hashing efficiency by adjusting parameters of the demultiplexing functions. For example, changing “positive.quantile” parameter in HTODemux function from a default value 0.99 to 0.9, increased hashing efficiency of the TotalSeq antibody nuclei dataset by more than 10% (from 54% to 65%) (**Sup. Fig. 6**). We also observed that a large number of negatives in MCF7 cluster (**Figure 7B**) is rather explained by an issue with the antibodies “hashtag 1”, since when other cells (DU145) were stained with antibodies from the same vial in a repeated experiment, many DU145 cells were also recognized as “Negatives” by Seurat (**Sup. Fig. 11B**). It is also worth mentioning that in this repeated “TotalSeq A nuclei rep2” experiment we reduced the number of captured nuclei to 9868 (there were 23451 nuclei in the first experiment). This potentially improved the hashing efficiency, which resulted in a higher number of correctly annotated singlets using MULTISeqDemux (63 % versus 50% in the first “TotalSeq A nuclei” experiment) (**Sup. Fig. 2**). Importantly, CMO nuclei hashing still outperformed the TotalSeq A nuclei hashing (85 % vs 77 %), when hashtag 1 (DU145) was excluded from the hashing efficiency calculation and compared to CMO nuclei hashing on three cell lines.

**Figure 6.**
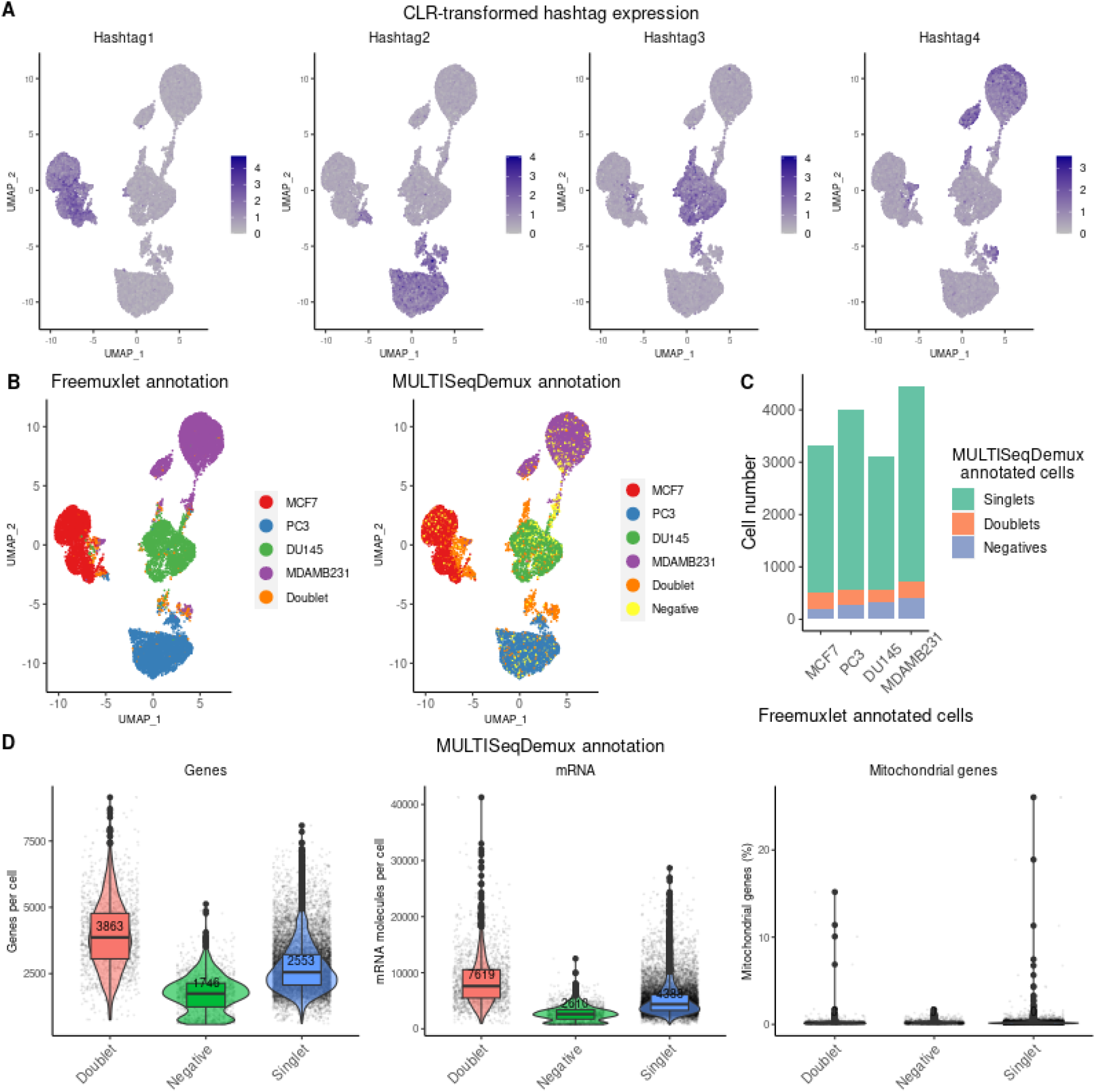
Cholesterol-modified oligo (CMO) hashing on nuclei. **A.** Gene-nuclei matrices were generated using CellRanger v3, followed by log-transformation of gene UMI counts and nuclei clustering (gene expression, PCA reduction) using Seurat. The hashtag UMI counts were CLR-transformed and visualised in blue color on the gene expression UMAP plots. **B.** Nuclei annotation (4 cell lines) was performed using freemuxlet (gene expression) or Seurat (MULTISeqDemux function applied on hashtag counts data) and visualized on the gene expression UMAP plots. **C.** MULTISeqDemux-annotated singlets (MCF7, PC3, DU145 or MDAMB231) and negatives (cells with background expression for each hashtag) were matched with the freemuxlet-annotated nuclei (MCF7, PC3, DU145 or MDAMB231). The rest (unmatched) of freemuxlet-annotated singlets were assigned as doublets and altogether visualized in barplots. **D.** Gene expression were log-transformed using Seurat and detected genes (left plot), UMIs (middle plot) and percentage of mitochondrial genes expression (right plot) in nuclei were visualised as violin-box plots with median values highlighted in red, across MULTISeqDemux-annotated groups (singlets, doublets, negatives on basis of hashtag expression).

**Figure 7.**
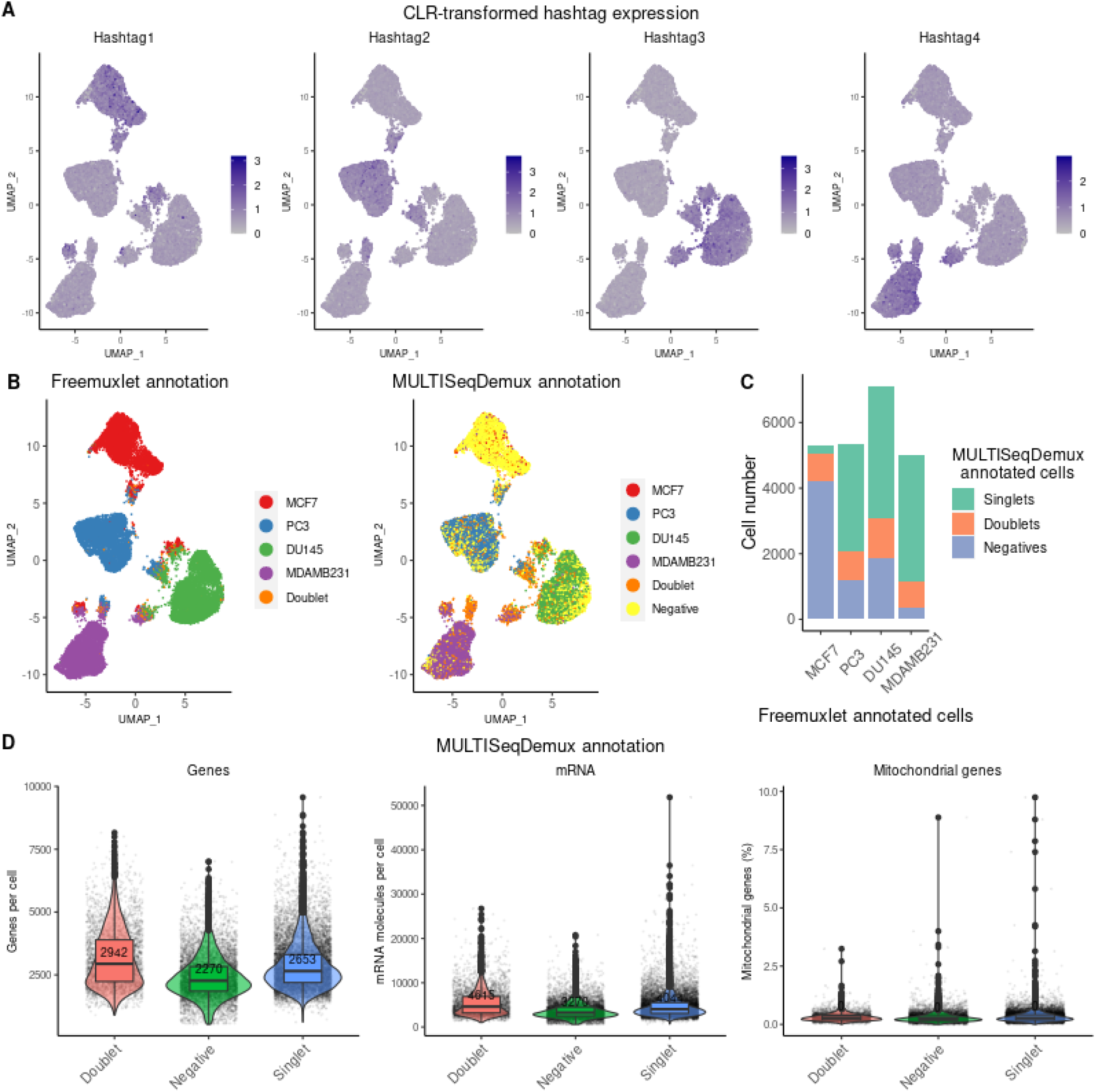
TotalSeq-A hashing on nuclei. **A.** Gene-nuclei matrices were generated using CellRanger, followed by log-transformation of gene UMI counts and nuclei clustering (gene expression, PCA reduction) using Seurat. The hashtag UMI counts were CLR-transformed and visualised in blue color on the gene expression UMAP plots. **B.** Nuclei annotation (4 cell lines) was performed using freemuxlet (gene expression) or Seurat (MULTISeqDemux function applied on hashtag counts data) and visualized on the gene expression UMAP plots. **C.** MULTISeqDemux-annotated singlets (MCF7, PC3, DU145 or MDAMB231) and negatives (cells with background expression for each hashtag) were matched with the freemuxlet-annotated nuclei (MCF7, PC3, DU145 or MDAMB231). The rest (unmatched) of freemuxlet-annotated singlets were assigned as doublets and altogether visualized in barplots. **D.** Gene expression were log-transformed using Seurat and detected genes (left plot), UMIs (middle plot) and percentage of mitochondrial genes expression (right plot) in nuclei were visualised as violin-box plots with median values highlighted in red, across MULTISeqDemux-annotated groups (singlets, doublets, negatives on basis of hashtag expression).

In the first experiment with more nuclei the antibody hashing efficiency, excluding the MCF7 cells (hashtag 1), only reached 64 %.

Interestingly, the tested hashing techniques on nuclei were characterised by an opposite pattern compared to cell hashing in respect to the background expression for each hashtag (reflected in number of detected “Negatives”). Accordingly, the MULTISeqDemux function annotated more droplets (33%) as “Negatives” in antibody nuclei hashing compared to CMO nuclei hashing experiment (8.1% droplets detected as negatives) (**Figure 5C** vs **6C**). This contrasts with the hashing of cells where antibody hashing delivered less “Negatives” compared to the lipid hashing (**Figure 2C, 3C** vs **4C, 5C**).

Additionally, we observed ~1,6 times less genes and UMIs per cell in “Negatives” compared to singlets in case of the lipid nuclei hashing (CMO) (**Figure 5D**). While for the antibody nuclei hashing experiment, this ratio was 1.2, for both number of UMIs and genes per cell (**Figure 6D**). The mislabelling hashtag rate for the nuclei samples varied from 0.89% of swapped labels among singlets for CMO technique to ~ 4% for TotalSeq-A nuclei hashing (**Sup. Table 4**).

We also wanted to compare some gene expression-related metrics in samples from the hashing experiments of this study and other non-hashed samples on the single cell line normalized to the same sequencing depth. We did not observe major differences in number of genes and UMIs per nucleus when comparing MCF7 nuclei from hashing (antibody and lipid) and non-hashed experiments (**Sup. Fig. 9**). The same conclusion was valid for all the tested hashing techniques on MCF7 cells (**Sup. Fig. 10**).

In general, opposite to the cell hashing we observed better hashing results on nuclei when using lipid-based hashing (CMO) compared to the antibody (TotalSeq-A) nuclei hashing.

### Antibody hashing can be successfully applied on clinical PBMC samples

In the context of the COVID-19 virus pandemic, we also tested TotalSeq-A antibody hashing on PBMCs from healthy individuals and SARS-CoV-2 patients in the frame of COVID-19 clinical study (NCT04326920). All samples additionally contained a large 277 antibody CITE-seq (TotalSeq-A antibodies) panel, thus making this PBMC hashing evaluation even more relevant for other multiomics scRNA-seq experiments. We could correctly assign 84% of all singlets to an appropriate patient with MULTISeqDemux (autoTresh = T) function and using freemuxlet annotation as a reference, in both pools: 3 healthy and 3 SARS-CoV-2 patients (**Table 2**). Cell clustering analysis demonstrated a presence of all major cell types (T cells, B cells, monocytes etc.) across 3 pooled samples in both pools (data not shown).

The MULTISeqDemux function annotated 5.2% and 6.3% cells as “Negatives” in antibody hashing of healthy and diseased samples, respectively (**Figure 8C, 9C**). The number of genes per cell in “Negatives” compared to singlets was 2.6 times lower (for UMI −3.3 times less) in a healthy control sample (**Figure 8D**). For the antibody hashing of diseased patients this ratio was 1.2 times lower for both number of UMIs and genes per cell (**Figure 9D**). Despite this, as we mentioned above, the hashing efficiency for both groups of PBMC samples was 84% (**Table 2**). According to freemuxlet, the hashtag mislabelling rate was 4.76% of swapped singlet labels for healthy sample and 1.53% for SARS-CoV-2 sample (**Sup. Table 4**).

**Figure 8.**
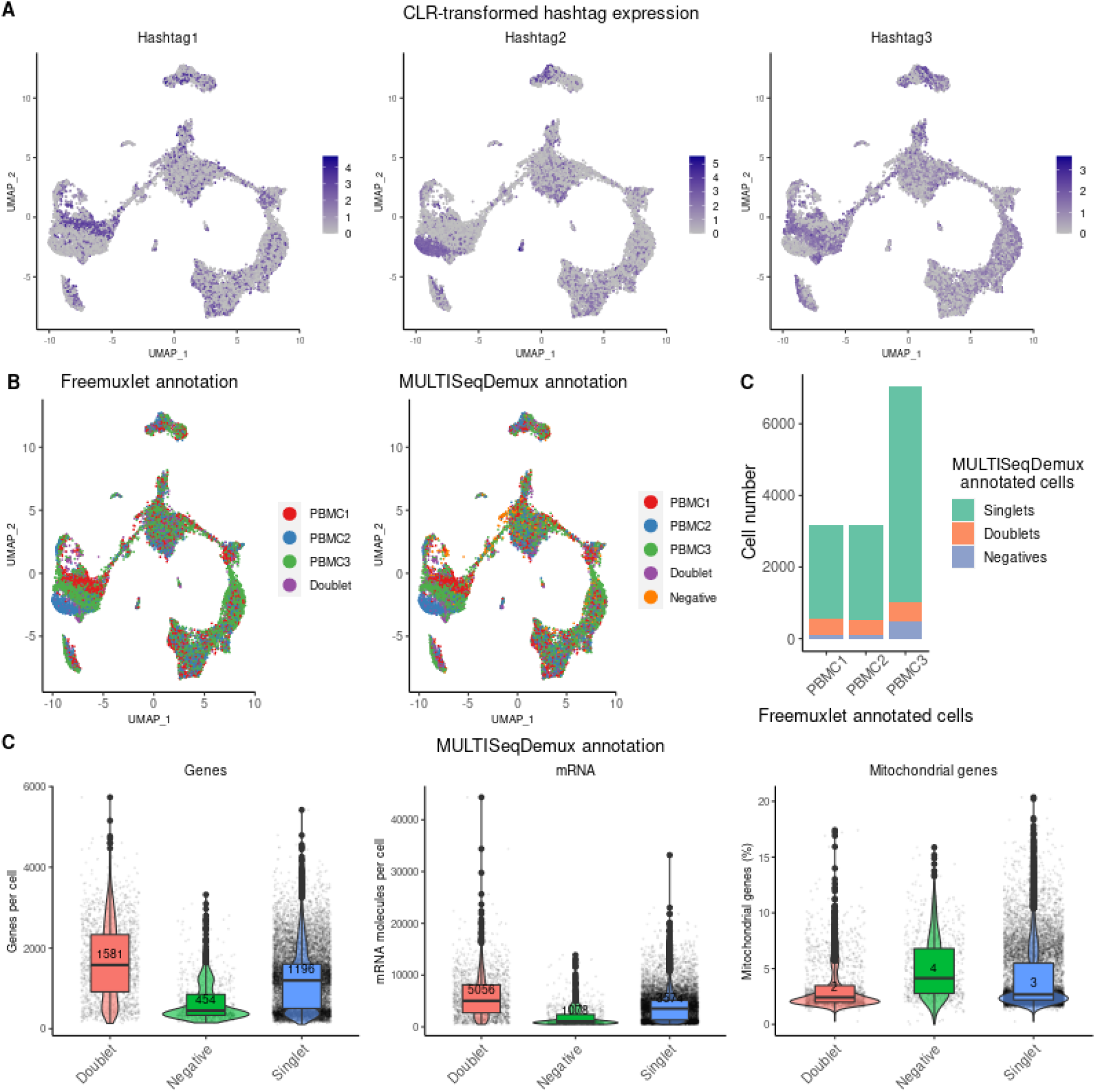
TotalSeq-A hashimg on PBMCs (healthy control). **A.** Gene-cell matrices were generated using CellRanger, followed by log-transformation of gene UMI counts and cell clustering (gene expression, PCA reduction) using Seurat. The hashtag UMI counts were CLR-transformed and visualised in blue color on the gene expression UMAP plots. **B.** Cell annotation (3 patients) was performed using freemuxlet (gene expression) or Seurat (MULTISeqDemux function applied on hashtag counts data) and visualized on the gene expression UMAP plots. **C.** MULTISeqDemux-annotated singlets (PBMC1, PBMC2 or PBMC3) and negatives (cells with background expression for each hashtag) were matched with the freemuxlet-based annotation (PBMC1, PBMC2 or PBMC3). The rest (unmatched) of freemuxlet-annotated singlets were assigned as doublets and altogether visualized in barplots. **D.** Gene expression were log-transformed using Seurat and detected genes (left plot), UMIs (middle plot) and percentage of mitochondrial genes expression (right plot) in cells were visualised as violin-box plots with median values highlighted in red, across MULTISeqDemux-annotated groups (singlets, doublets, negatives on basis of hashtag expression).

**Figure 9.**
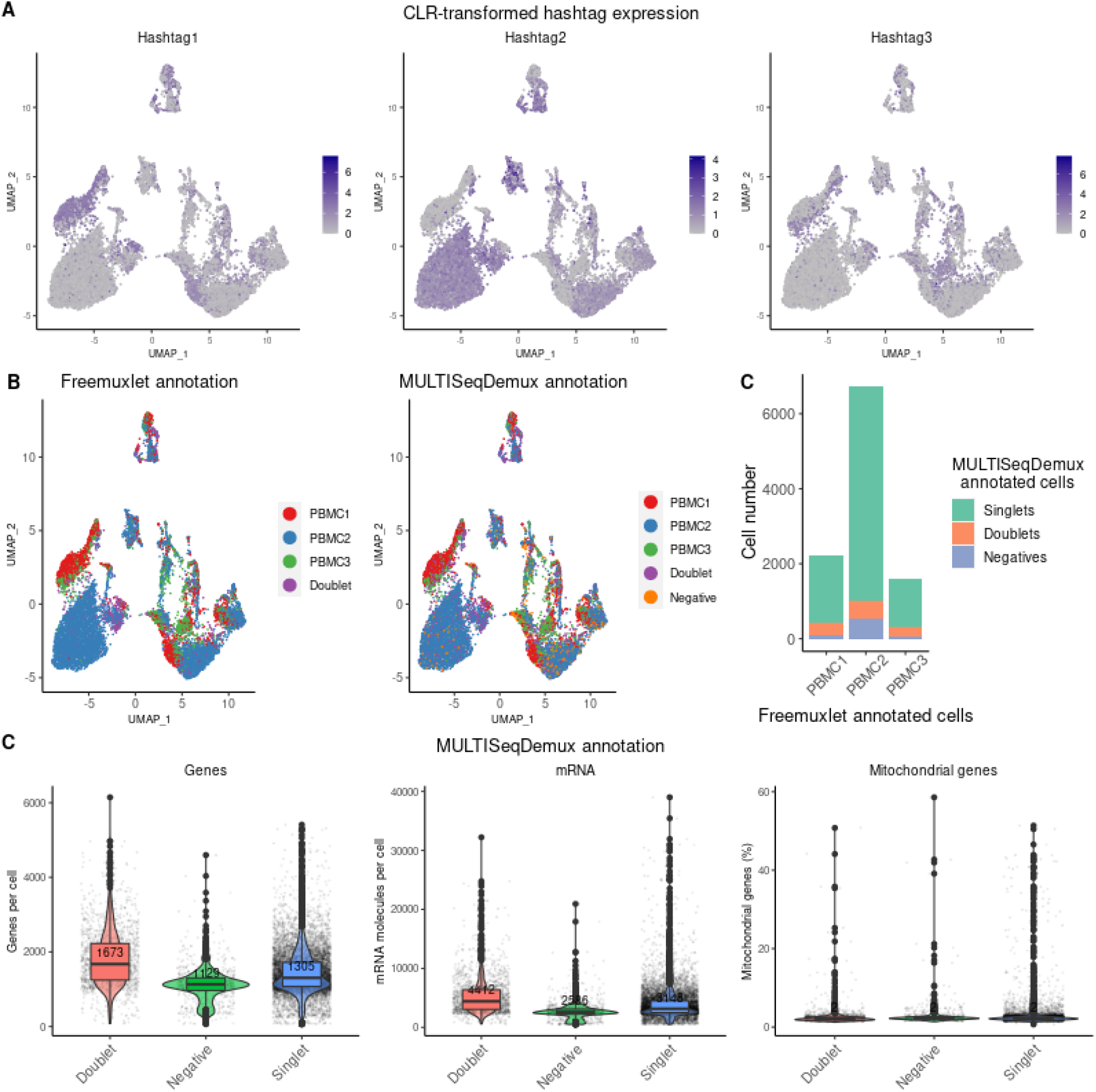
TotalSeq-A hashing on PBMCs (SARS-CoV-2 patients). **A.** Gene-cell matrices were generated using CellRanger, followed by log-transformation of gene UMI counts and cell clustering (gene expression, PCA reduction) using Seurat. The hashtag UMI counts were CLR-transformed and visualised in blue color on the gene expression UMAP plots. **B.** Cell annotation (3 patients) was performed using freemuxlet (gene expression) or Seurat (MULTISeqDemux function applied on hashtag counts data) and visualized on the gene expression UMAP plots. **C.** MULTISeqDemux-annotated singlets (PBMC1, PBMC2 or PBMC3) and negatives (cells with background expression for each hashtag) were matched with the freemuxlet-based annotation (PBMC1, PBMC2 or PBMC3). The rest (unmatched) of freemuxlet-annotated singlets were assigned as doublets and altogether visualized in barplots. **D.** Gene expression were log-transformed using Seurat and detected genes (left plot), UMIs (middle plot) and percentage of mitochondrial genes expression (right plot) in cells were visualised as violin-box plots with median values highlighted in red, across MULTISeqDemux-annotated groups (singlets, doublets, negatives on basis of hashtag expression).

**Figure 10.**
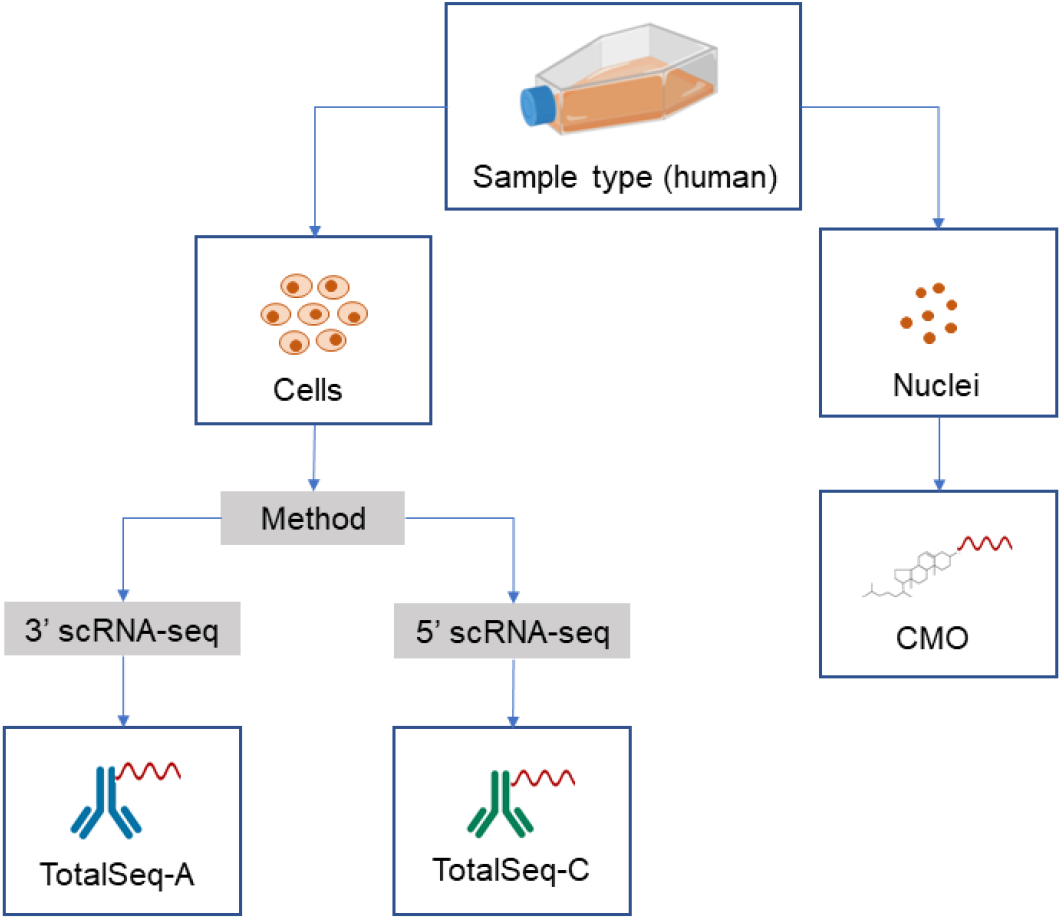
Preferable hashing techniques according to our study om human cancer cell lines (MCF7, PC3, DU145 and MDAMB231). TotalSeq antibody hashing is a preferred technique for multiplexing different human cell lines. The CMO hashing method yields the best results when nuclei are the input sample. CMO - cholesterol-modified oligos.

## DISCUSSION

Correct sample demultiplexing after hashing depends on several factors: the hashing strategy itself (antibodies versus lipids), which subsequently can depend on the cell surface antigen repertoire in case of antibody hashing or biochemical/biophysical properties of membranes in case of lipid hashing. Also, this accuracy depends on the demultiplexing methods that can be subsequently influenced by other parameters (cell number, cells or nuclei, sequencing depth etc.). In order to accurately determine the efficiency of demultiplexing, we hashed different cell lines that carry intrinsic genetic variations which serve as a ground truth control. Importantly, for one of the hashing datasets we compared two reference demultiplexing tools: freemuxlet which was used in this study and genotyping data-dependent tool demuxlet (Kang et al. 2018). The cell deconvolution of “TotalSeq-A on cells” sample using HTODemux demultiplexing function demonstrated 94% singlet concordance with freemuxlet output as a reference, and 95% concordance with demuxlet as a reference. Our comparison demonstrates that even without external genotyping data (SNPs), required for demuxlet, it is possible to deconvolute different cells using a similar algorithm (freemuxlet). We noticed however that “doublet vs singlet” classification by freemuxlet was not always consistent with the annotation by DoubletFinder (Mcginnis et al. 2019) or Scrublet, the commonly used tools for computational doublet prediction (Wolock et al. 2019). For some data sets freemuxlet was underestimating the number of doublets. Additionally, the doublets annotated by HTO- and MULTISeqDemux were also not always aligned with the output of DoubletFinder and Scrublet. These findings are in accordance with the observation of a relatively high disagreement between DoubletFinder, Scrublet and demuxlet (DePasquale et al. 2019). This can be partially explained by the reliance on different data modalities that can lead to a certain bias in detection of heterotypic or homotypic doublets. For instance, the functions incorporated in Seurat predict doublets based on the hashtag signal, while other tools use the gene expression data. Overall, this limitation in correct doublet prediction might diminish demultiplexing accuracy and requires further optimisation. The impaired doublet annotation together with a failure to assign hashtags to a number of cells (“negatives”) might explain the lower hashing efficiency in the samples with higher number of cells. For example, the cell number in the repeated TotalSeq A experiment (rep2) was increased by 48%, while the number of doublets (detected by freemuxlet) and negatives (assigned by MULTISeqDemux) increased by 125% and 325%, respectively.

It is important to mention that our comparison of different hashing strategies was applied to cells from filtered gene-cell matrices as produced by CellRanger v3 without further outlier filtering (e.g. based on mitochondrial gene expression). We could see in our PBMC dataset that in terms of cell calling, this filtered matrix contains almost the same number of cells as a “raw” matrix after filtering out all droplets with less than 200 UMIs (gene expression) and all droplets expressing one gene in less than 3 droplets. We observed that the tested demultiplexing functions from Seurat assigned some cells with relatively low number of genes and UMIs to the group “negatives”. Hence, not surprisingly the hashing efficiency can be improved by filtering out the cell outliers. For example, correct PBMCs annotation (healthy control patients) was improved from 84,1% to 89,4 % after filtering out 2660 from 14630 cells using the “scater” Bioconductor package (outliers on basis of number of genes and UMIs per cell). On the other hand, a stringent filtering might cause a loss of biologically relevant cell subtypes. Thus, optimisation of the sample demultiplexing should be performed in parallel with a deeper biological sample analysis.

We can conclude that the type of hashing strategy must be chosen based on the hashtag antigen expression, since it is known that some cells might not express both antigens targeted by available TotalSeq-hashing antibodies. For instance, CD45 and MHC II antigens (targeted by mouse hashing antibodies from BioLegend) in mouse C3 and B16-BL6 melanoma cells (**Sup. Fig. 8**).

However, a relatively lower affinity of certain cells towards different type of lipid hashes can be also observed. Nevertheless, lipid-based hashing might be a preferred strategy for multiplexing of samples with a low or unknown expression of hashing antigens, provided the sample demultiplexing is well optimized.

In the present study on human cancer cell lines, the antibody-based hashing when targeting cells overall performed better than the lipid-based hashing. When labelling nuclei from the same cell lines, cholesterol-based hashes (CMO) demonstrated better hashing efficiency compared to the antibody-based labelling (TotalSeq -A). This result can be partially explained by the potential detrimental effect of the lysis buffer (Nuclei EZ Lysis Buffer) used for nuclei isolation on the nuclei surface proteins. The comparison of TotalSeq -A based cell hashing with TotalSeq-C cell hashing compatible with 3’ and 5’ gene 10x Genomics sequencing respectively, demonstrates similarly good hashing efficiency. Alternatively, other antibody-based hashing methods are also available on the market, e.g. hashing antibodies from BD Biosciences compatible with the BD Rhapsody platform for scRNA-seq experiments.

With respect to analysis algorithms, we observed that the MULTISeqDemux function with autoTresh =T parameter originally tailored for lipid-based hashing (McGinnis et al. 2019), performs even better than HTODemux function on a vast majority of tested data sets.

Overall, we can conclude that various hashing techniques can be successfully applied on cells and nuclei, with different hashing efficiency metrics though, to reduce costs and batch effects of scRNA-seq experiments.

## Acknowledgements

We gratefully acknowledge the help of Gert Van Isterdael with flow cytometry experiments and the analysis, Christopher McGInnis from UCSF for providing the MULTI-seq reagents, Mark Fiers for providing with pre-mRNA genome, Kris Davie for providing us with the vcf files, Kevin Verstaen for help with optimising CellRanger and for valuable discussions on analysis, and Junbin Qian for optimising and running demuxlet. This work was funded by the “VIB Tech Watch – Janssen” collaboration.

## Author contributions

V.M. performed experiments, did analysis, wrote the manuscript. J.A. performed experiments, wrote the manuscript. I.M. did analysis, wrote the manuscript. S.P., N.V. performed experiments. G.H. did analysis. Y.S., S.A., S.VH., S.P., N.V., H.W., J.R., J.VH., N.V., R.S. supervised the work and reviewed the manuscript.

## Disclosure statement

No potential conflict of interest was reported by the authors.

## Notes

### Competing Interest Statement

The authors have declared no competing interest.

