## Supplementary material for "Comparative analysis of antibody- and lipid-based multiplexing methods for single-cell RNA-seq": Sup. Fig. 1

### SUPPLEMENTAL DATA

**A**

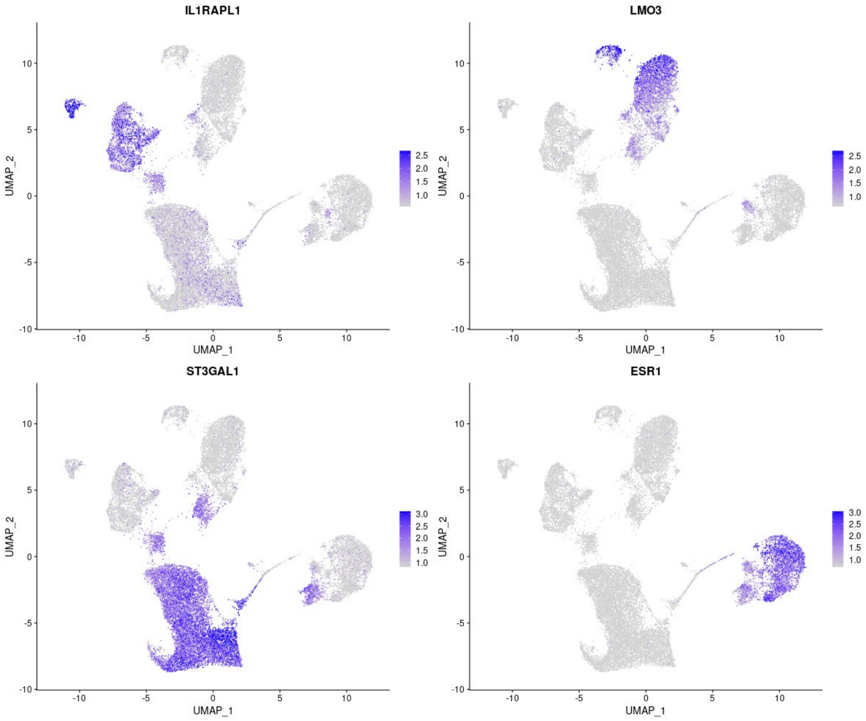


**B**

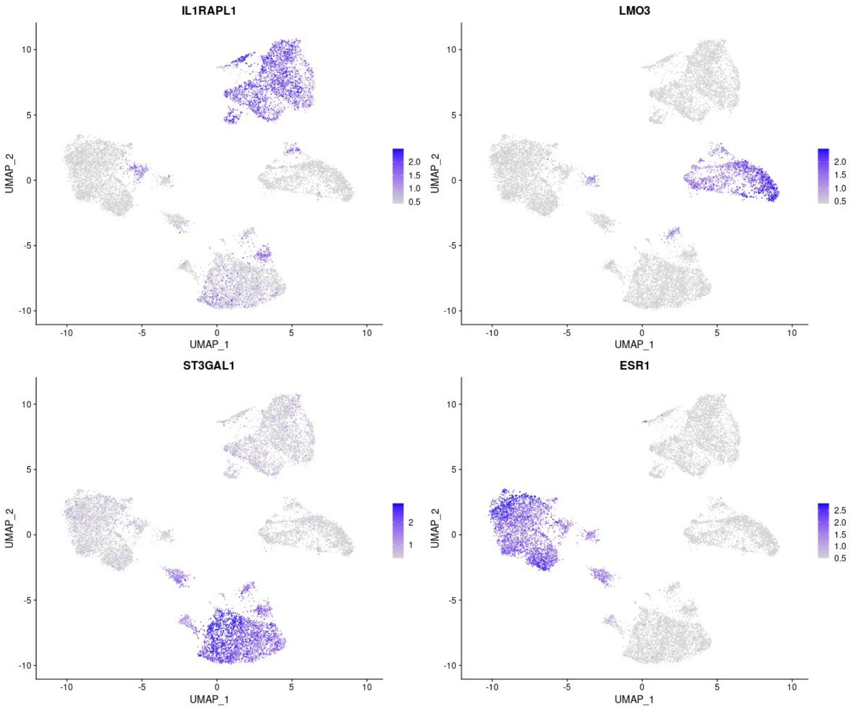


**Sup. Fig. 1. Marker gene expression in TotalSeq-A (A) and custom lipid (B) cell hashing samples.** Gene-cell matrices were generated using CellRanger, followed by log-transformation of gene UMI counts and cell clustering (gene expression, PCA reduction) using Seurat. The marker gene UMI counts were visualised in blue color on the gene expression UMAP plots.

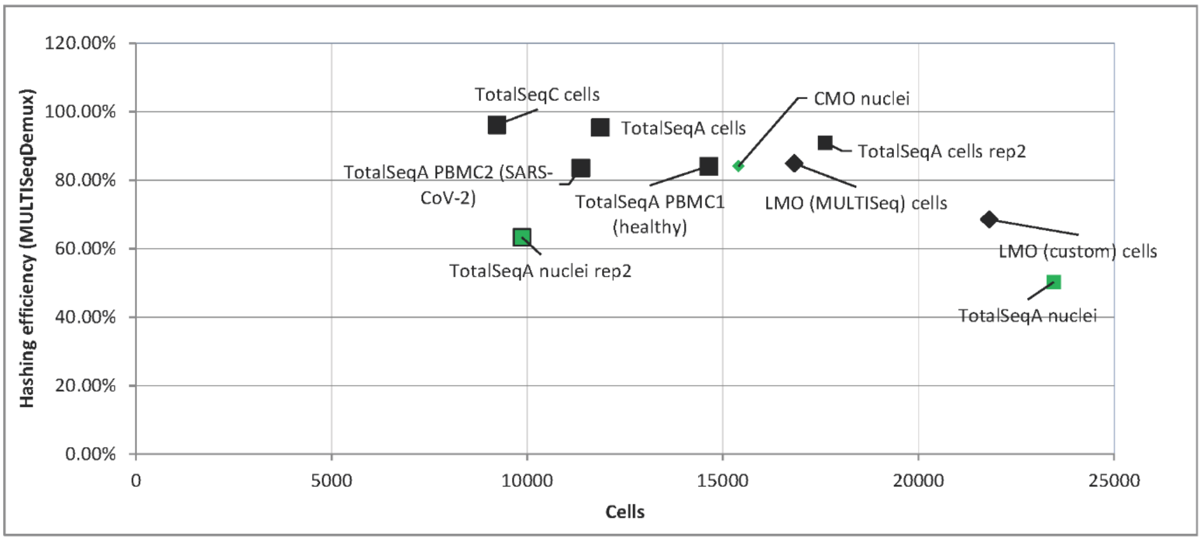


**Sup. Fig. 2. Dependency between cell number and hashing efficiency.** The hashing efficiency was calculated as an overlap between cell line annotation using Seurat (MULTISeqDemux) and freemuxlet as a reference annotation. Black color – cell hashing; green – nuclei hashing; square – hashing using antibodies; rhombus – hashing using lipids. The plot created using an excel macro: Copyright © 2009, Clear Lines Consulting, LLC.

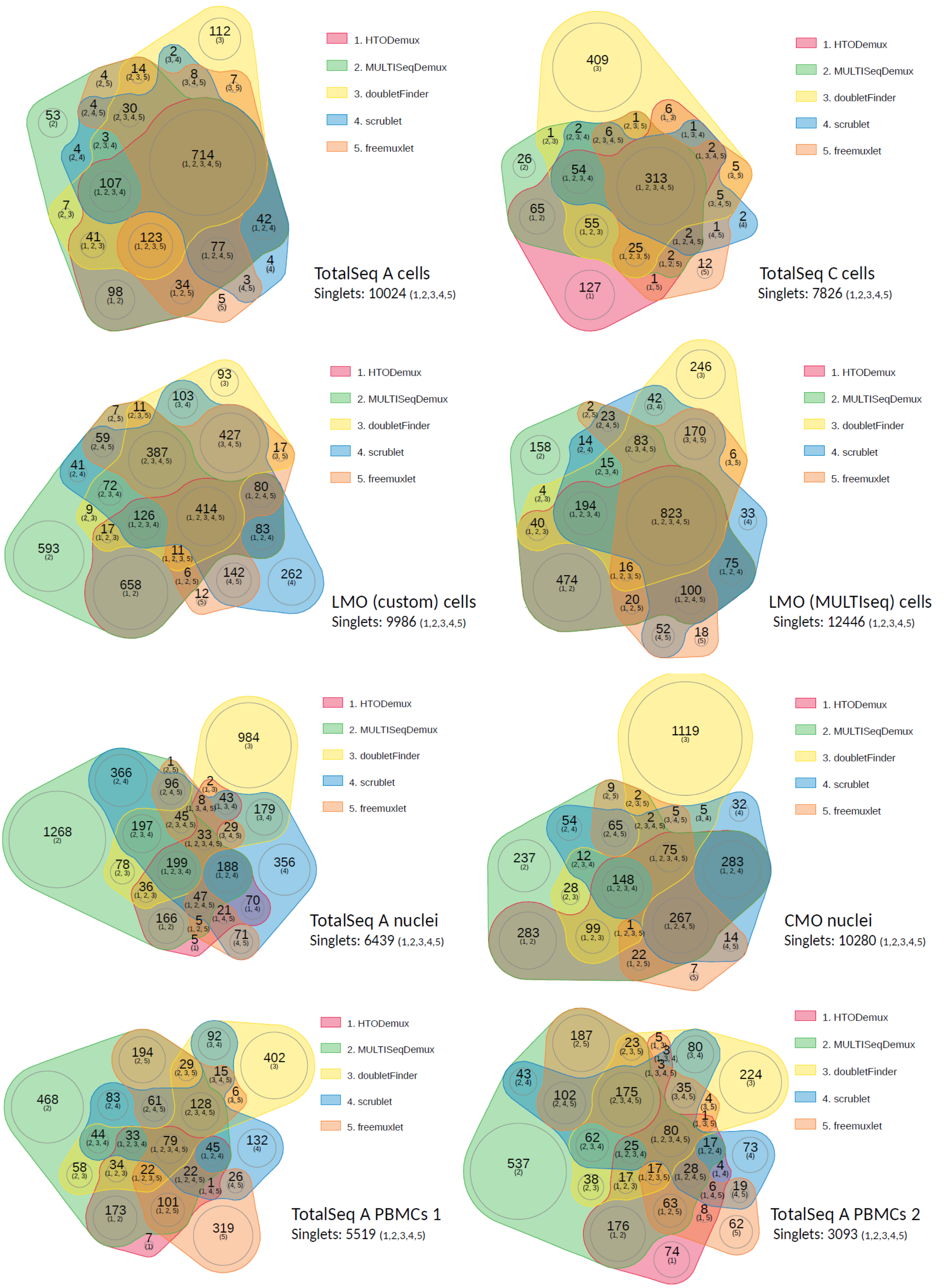


**Sup. Fig. 3. Number of detected doublets by different doublet annotation tools depicted as venndiagrams** ("nVennR" package). In parenthesis – which tools compared. For a comparison, number of singlets detected by all 5 tools (1,2,3,4,5) is also shown for each experiment (in the corner).

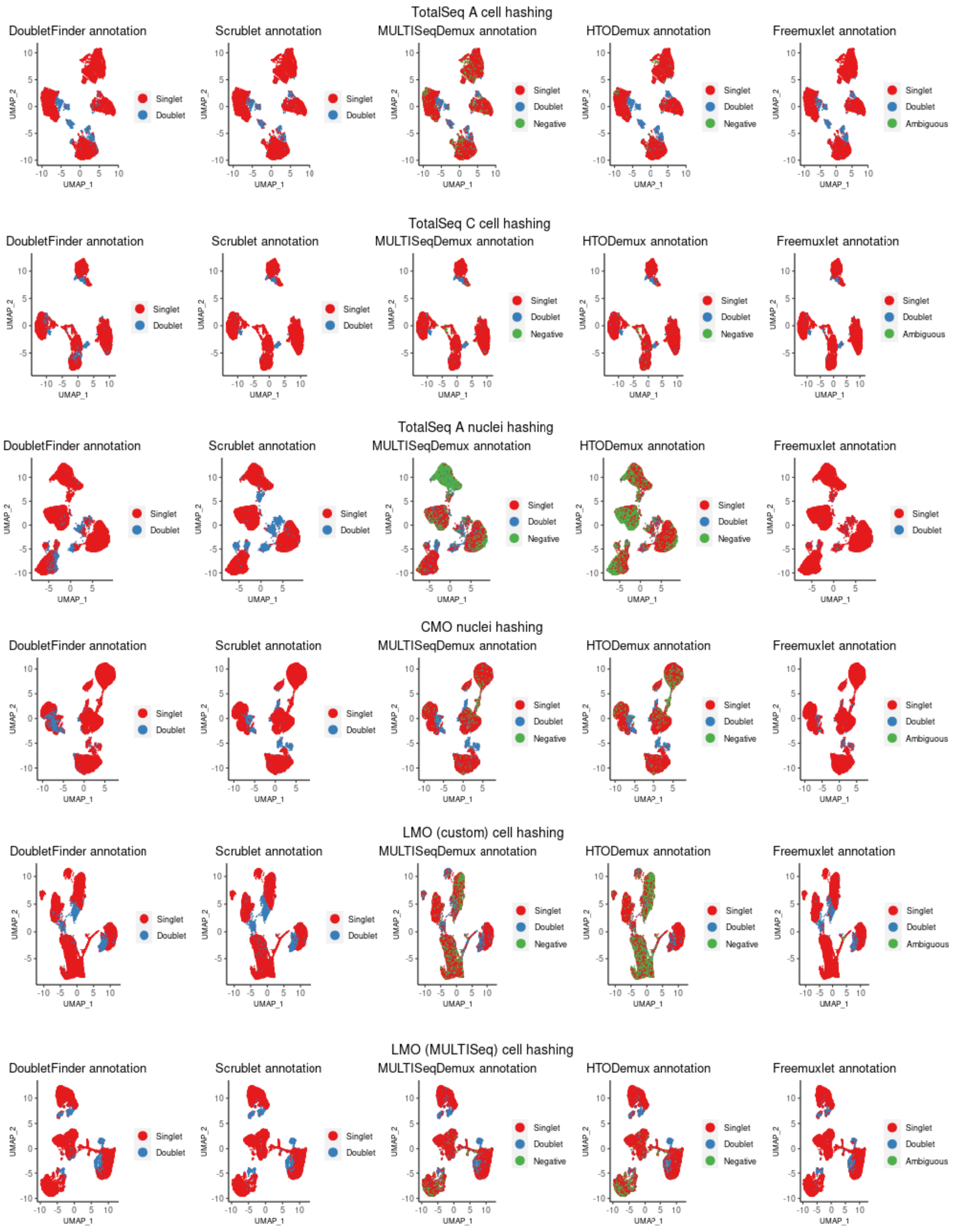


**Sup. Fig. 4. Comparison of doublet annotations (4 cancer cell lines).** Gene-cell matrices were generated using CellRanger v3, followed by log-transformation of gene UMI counts and cell clustering (gene expression, PCA reduction) using Seurat. Droplet annotation using 5 different methods is depicted on the gene expression UMAP plots. MULTISeqDemux and HTODemux are the functions from Seurat.

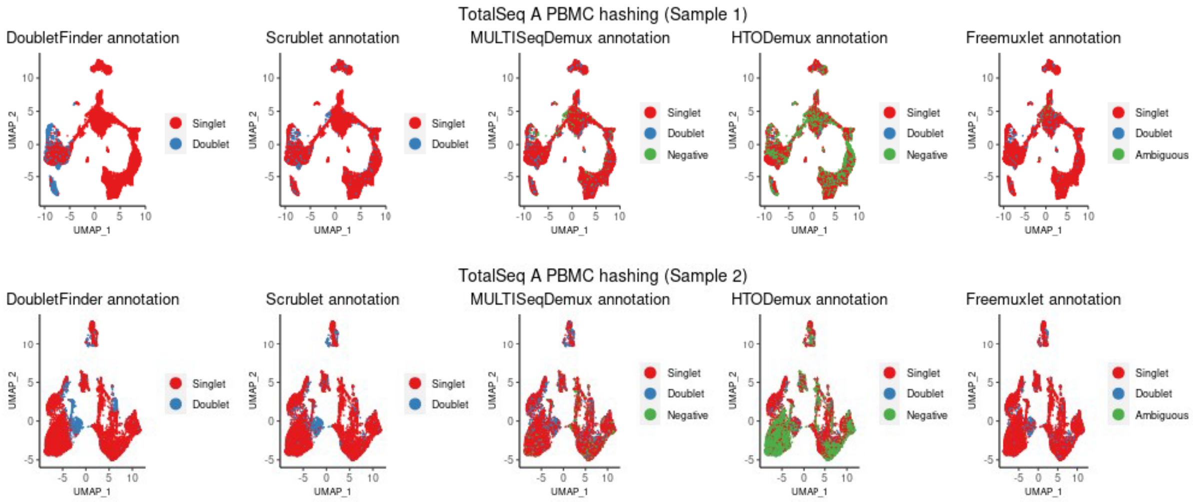


**Sup. Fig. 5. Comparison of doublet annotations (PBMCs).** Gene-cell matrices were generated using CellRanger v3, followed by log-transformation of gene UMI counts and cell clustering (gene expression, PCA reduction) using Seurat. Droplet annotation using 5 different methods is depicted on the gene expression UMAP plots. MULTISeqDemux and HTODemux are the functions from Seurat. Sample 1 – healthy patients. Sample 2 – COVID-19 patients.

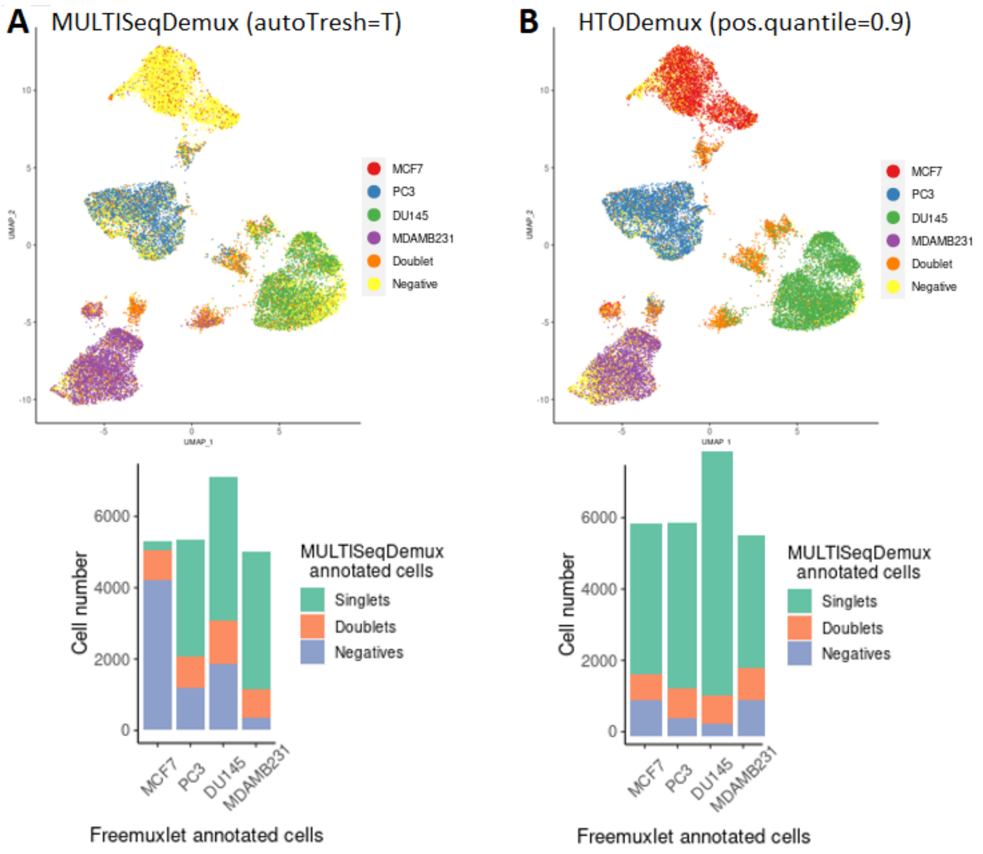


**Sup. Fig. 6. Finetuning demultiplexing.** MULTISeqDemux (autoTresh=T) annotation (**A**) vs HTODemux (pos.quant.=0,9) (**B**) on TotalSeq-A nuclei hashing sample. Nuclei annotation (4 cell lines) was performed using freemuxlet (gene expression) or Seurat (MULTISeqDemux function applied on hashtag counts data) and visualized on the gene expression UMAP plots. For the barplots above, MULTISeqDemux-annotated singlets (MCF7, PC3, DU145 or MDAMB231) and negatives (cells with background expression for each hashtag) were matched with the freemuxlet-based annotation (MCF7, PC3, DU145 or MDAMB231). The rest (unmatched) of freemuxlet-annotated singlets were assigned as doublets and altogether visualized as barplots.

**A**

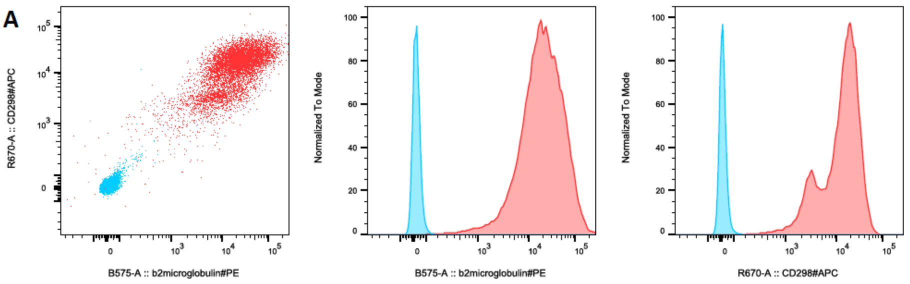


| Sample Name | Subset Name | Count | PE-pos :: Freq. of Parent | APC-pos :: Freq. of Parent |
|---|---|---|---|---|
| PBMCs | cells | 40549 | 96.5% | 98.2% |
| no abs PBMCs | cells | 39514 | 5.06E-3% | 0.020% |

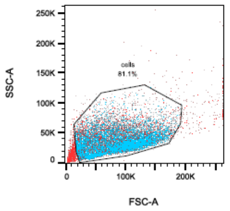


**B**

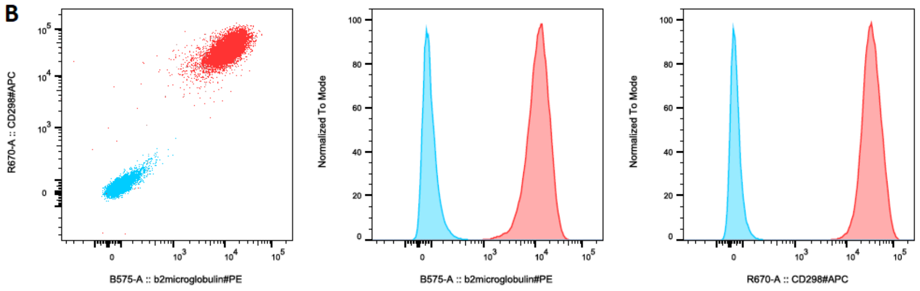


| Sample Name | Subset Name | Count | PE-pos :: Freq. of Parent | APC-pos :: Freq. of Parent |
|---|---|---|---|---|
| HEK293A | cells | 37391 | 98.8% | 99.9% |
| no abs HEK293A | cells | 38519 | 0% | 0% |

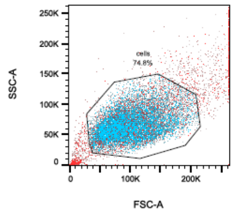


**C**

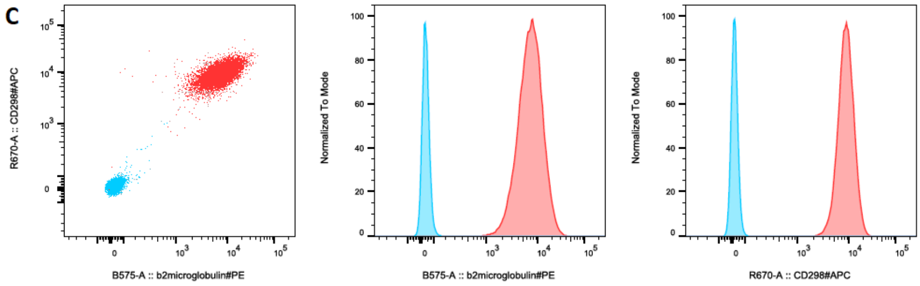


| Sample Name | Subset Name | Count | PE-pos :: Freq. of Parent | APC-pos :: Freq. of Parent |
|---|---|---|---|---|
| THP-1 | cells | 35441 | 97.9% | 100.0% |
| no abs THP-1 | cells | 37489 | 0% | 0.032% |

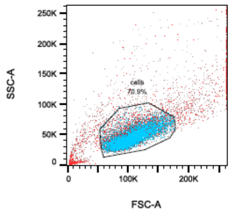


**Sup. Fig. 7.** Expression of hashing antigens on human PBMCs (**A**), HEK293A (**B**) and THP-1 (**C**) cells detected by flow cytometry using the same CD298 and b2-microglobulin clones as in the human hashing TotalSeq antibodies. Red color – cells with the antibody staining; blue color – cells without the staining (negative control). Other cells that express both antigens: HeLa, Jurkat, 501-mel, MDA-MB-231, A375m, HIBCPP, HaCaT, BLM, OVCAR-3, HT-29, human fibrosarcoma cells, ARPI9, CAOV-3, HCT116.

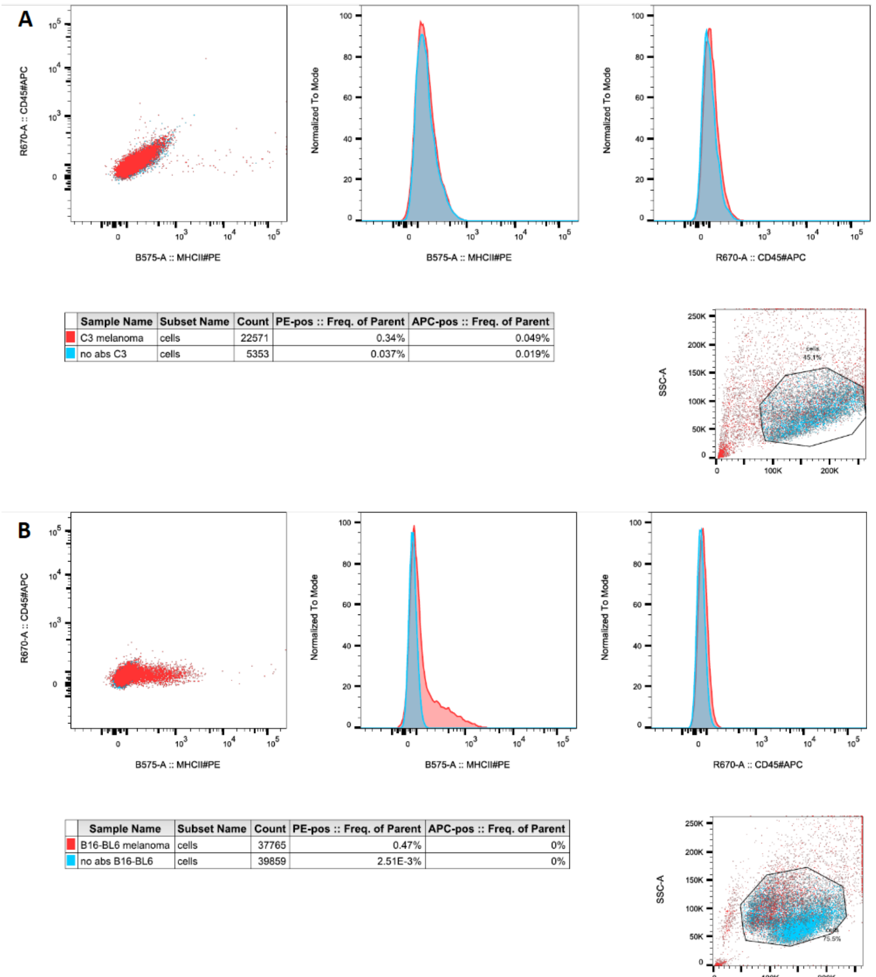


| Sample Name | Subset Name | Count | PE-pos :: Freq. of Parent | APC-pos :: Freq. of Parent |
|---|---|---|---|---|
| C3 melanoma | cells | 22571 | 0.34% | 0.049% |
| no abs C3 | cells | 5353 | 0.037% | 0.019% |

| Sample Name | Subset Name | Count | PE-pos :: Freq. of Parent | APC-pos :: Freq. of Parent |
|---|---|---|---|---|
| B16-BL6 melanoma | cells | 37765 | 0.47% | 0% |
| no abs B16-BL6 | cells | 39859 | 2.51E-3% | 0% |

**Sup. Fig. 8.** Expression of hashing antigens (CD45 and MHC II) on mouse C3 (**A**) and BL6 melanoma (**B**) cells detected by flow cytometry (same antibody clones as in mouse hashing TotalSeq antibodies). Red color – cells with the antibody staining; blue color – cells without the staining (negative control). The only mouse cell type that expressed both antigens (CD45 and MHC II) from the 8 tested mouse cell lines was J774A1 (macrophages).

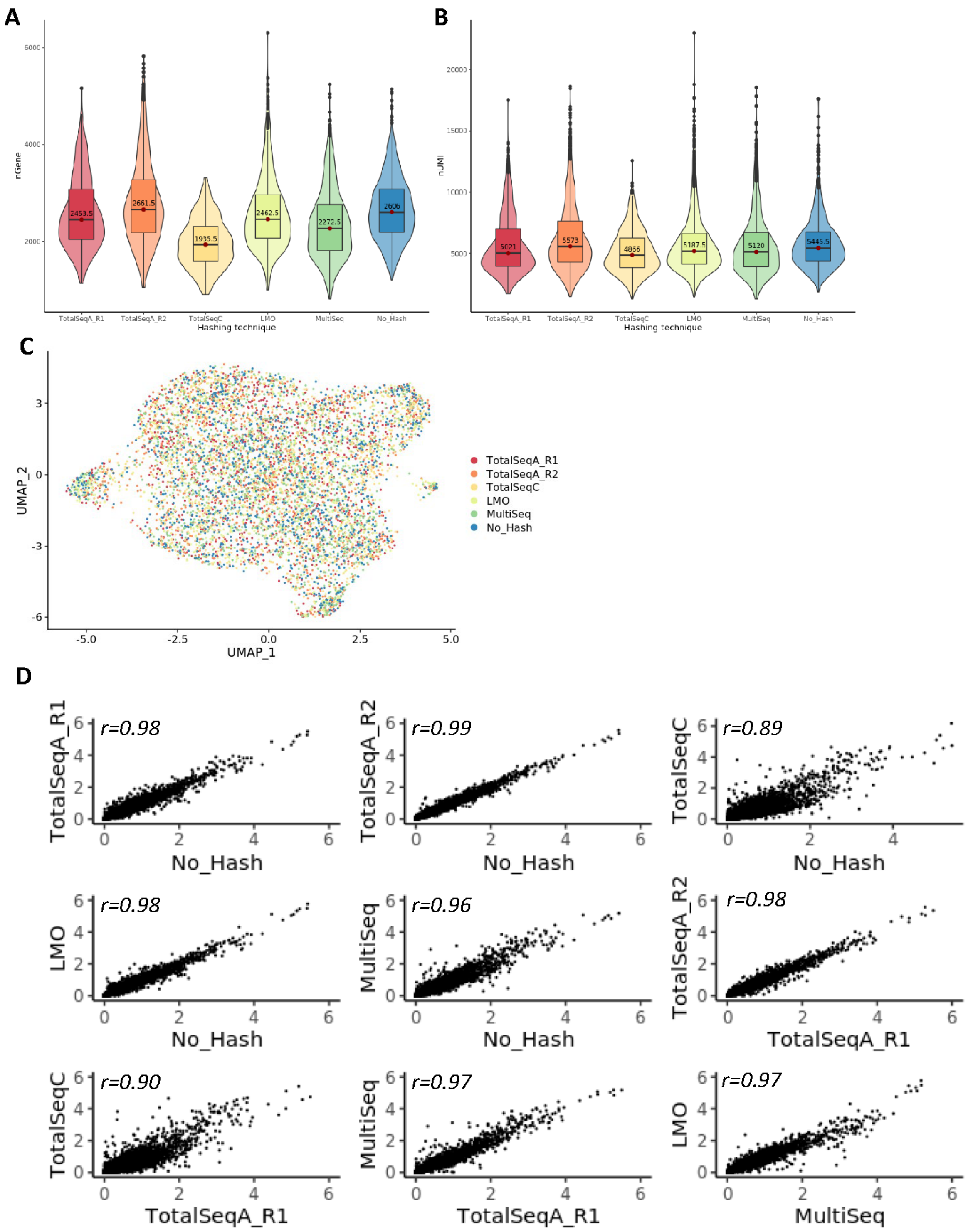


**Sup. Fig. 9. Similarity measures of the gene expression on MCF7 cells across the hashing strategies.**

Gene-cell matrices were generated using CellRanger v.3.1, followed by log-transformation of gene UMI counts and cell clustering (gene expression, PCA reduction) using Seurat. The hashtag UMI counts were CLR-transformed. To correct the effect of differences in sequencing depth per sample, the reads for each method were down-sampled (DropletUtilis), to match the protocol with the lowest number of total reads (TotalSeq-C). To the hashtag demultiplexed Seurat objects freemuxlet and DoubletFinder results were added as metadata. For each hashed sample the MCF7 cells were extracted from the Seurat object by filtering for i) cells with a valid MCF7 hashtag; ii) cells assigned by freemuxlet as MCF7 genotype and iii) cells assigned as singlets by DoubletFinder. The unhashed sample was processed in the same manner except for filtering for MCF7 hashtags. Each MCF7 cells subset was downsized to the hashing method with lowest number of MCF7 cells (TotalSeq-C 1178 cells) and filtered for low quality cells (low number of genes per cell and high % of mitochondrial genes). Next, the gene expression across the different hashing technologies were normalized, cell cycle and mitochondrial genes were regressed and data were integrated using the SCTransform workflow and **A** detected genes, **B** UMIs in MCF7 cells were visualised as violin-box plots with median values highlighted. **C**. The UMAP visualizations on the integrated gene expression. Cells are colored by hashing technology. **D.** Average log (gene expression) scatter plots across the hashing technologies on down-sampled data (5000), r= Pearson'scorrelation.

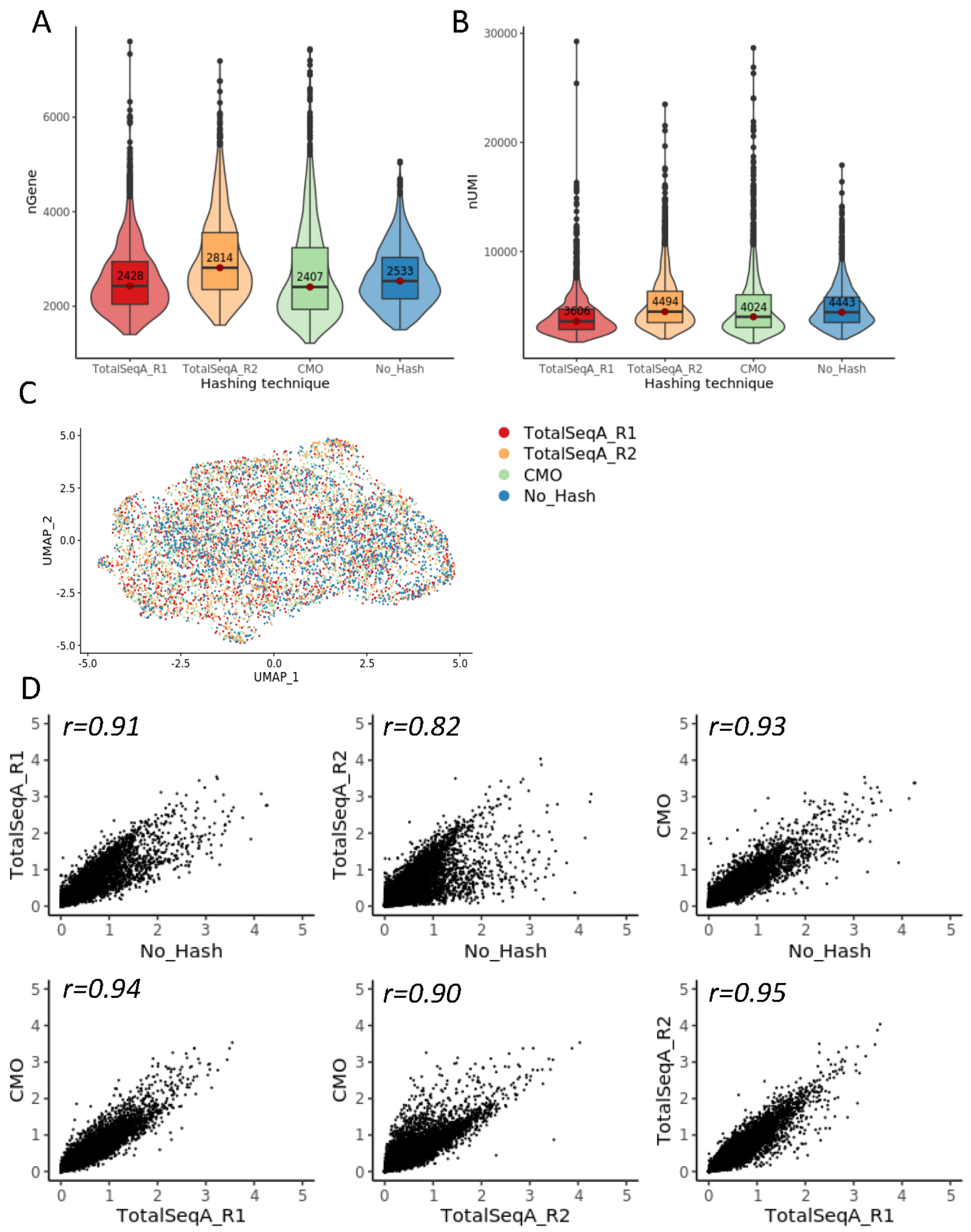


**Sup. Fig. 10. Similarity measures of the gene expression on MCF7 nuclei across the hashing strategies.**

Gene-nuclei matrices were generated using CellRanger v.3.1, followed by log-transformation of gene UMI counts and nuclei clustering (gene expression, PCA reduction) using Seurat. The hashtag UMI counts were CLR-transformed. To correct the effect of differences in sequencing depth per sample, the reads for each method were down-sampled (DropletUtilis), to match the protocol with the lowest number of total reads (TotalSeq-A_rep2). To the hashtags demultiplexed Seurat objects freemuxlet and DoubletFinder results were added as metadata. For each hashed sample the MCF7 nuclei were extracted from the Seurat object by filtering for i) nuclei with a valid MCF7 hashtag; ii) nuclei assigned by freemuxlet as MCF7 genotype and iii) nuclei assigned as singlets by DoubletFinder. The unhashed sample was processed in the same manner except for filtering for MCF7 hashtags. Each MCF7 nuclei subset was downsized to the hashing method with lowest number of MCF7 nuclei (TotalSeq-A_rep2 2726 cells) and filtered for low quality nuclei (low number of genes per nucleus and high % of mitochondrial genes). Next, the gene expression across the different hashing technologies were normalized, cell cycle and mitochondrial genes were regressed and data were integrated using the SCTransform workflow and **A** detected genes, **B** UMIs in MCF7 nuclei were visualised as violin-box plots with median values highlighted. **C**. The UMAP visualizations on the integrated gene expression. Nuclei are colored by hashing technology. **D.** Average log (gene expression) scatter plots across the hashing technologies on down-sampled data, r= Pearson's correlation.

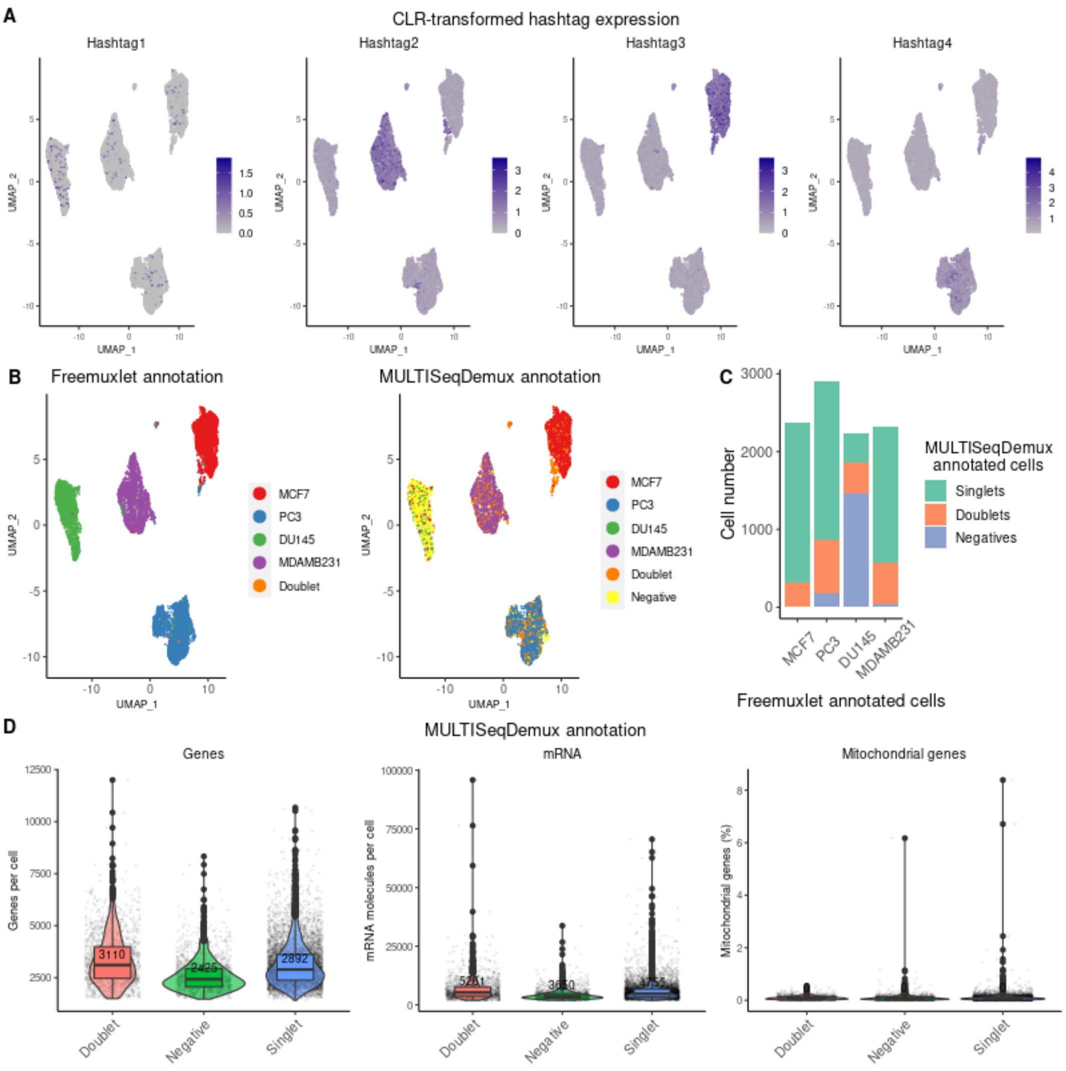


**Sup. Fig. 11. TotalSeq-A hashing on nuclei (a repeat experiment with less nuclei and reshuffled hashing antibodies)**. **A**. Gene-nuclei matrices were generated using CellRanger, followed by log-transformation of gene UMI counts and nuclei clustering (gene expression, PCA reduction) using Seurat. The hashtag UMI counts were CLR-transformed and visualised in blue color on the gene expression UMAP plots. **B**. Nuclei annotation (4 cell lines) was performed using freemuxlet (gene expression) or Seurat (MULTISeqDemux function applied on hashtag counts data) and visualized on the gene expression UMAP plots. **C**. MULTISeqDemux-annotated singlets (MCF7, PC3, DU145 or MDAMB231) and negatives (cells with background expression for each hashtag) were matched with the freemuxlet-based annotation (MCF7, PC3, DU145 or MDAMB231). The rest (unmatched) of freemuxlet-annotated singlets were assigned as doublets and altogether visualized as barplots. **D**. Gene expression were log-transformed using Seurat and detected genes (left plot), UMIs (middle plot) and percentage of mitochondrial genes expression (right plot) in nuclei were visualised as violin-box plots with median values highlighted in red, across MULTISeqDemux-annotated groups (singlets, doublets, negatives on basis of hashtag expression).

**Sup. Table 1**. **TotalSeq-A anti-nucleoporin antibody barcode sequences:**

| Hashtag | Antibody barcode |
|---|---|
| 1. | A0458 TGACGCCGTTGTTGT |
| 2. | A0459 GCCTAGTATGATCCA |
| 3. | A0456 CTCGAACGCTTATCG |
| 4. | A0457 CTTATCACCGCTCAA |

**Sup. Table 2. LMOs, CMOs and sample barcode oligonucleotides:**

| | |
|---|---|
| Custom LMO Anchor: | 5’-AGTGACAGCTGGATCGTTAC[Palmitate]-3’ |
| Custom LMO Co-anchor: | 5’-[Stearyl]GTAACGATCCAGCTGTCACTCACGTCTGAACTCCAGTCAC-3’ |
| CMO Anchor: | 5’AGTGACAGCTGGATCGTTAC[Chol-TEG]-3’ |
| LMO Co-anchor: | 5’[Chol-TEG]GTAACGATCCAGCTGTCACTCACGTCTGAACTCCAGTCAC-3’ |
| Hashtag 1 | TTGTCACGGTAATTA |
| Hashtag 2 | ATCGAACCGACAGAG |
| Hashtag 3 | GGTCGAATATGTCGG |
| Hashtag 4 | CTCAAGCATTATCAT |

**Sup. Table 3**. **Comparison of doublets detected by freemuxlet and negatives detected by MULTISeqDemux:**

| Experiment | Estimated Number of Cells | Hashing efficiency (MULTISeq Demux) | Number of doublets (freemuxlet) | Number of MULTISeq-annotated singlets among freemuxlet doublets | % of MULTISeq-annotated singlets among freemuxlet doublets | Number of MULTISeq-annotated negatives |
|---|---|---|---|---|---|---|
| 1. TotalSeq-A cells | 11869 | 95.4 % | 1031 | 25 | 2.42 % | 132 |
| 2. TotalSeq-A cells rep2 | 17611 | 90.9 % | 2325 | 111 | 4.77 % | 516 |
| 3. TotalSeq-C cells | 9229 | 96.2 % | 413 | 26 | 6.29 % | 165 |
| 4. LMO (MULTI-seq) cells | 16827 | 84.9 % | 1326 | 253 | 19.07 % | 1230 |
| 5. LMO (custom) cells | 21813 | 68.5 % | 2052 | 777 | 37.86 % | 4542 |
| 6. CMO nuclei | 15404 | 84.1 % | 543 | 38 | 6.99 % | 1253 |
| 7. TotalSeq-A nuclei | 23451 | 50.2 % | 550 | 182 | 33.09 % | 7729 |
| 8. TotalSeq-A nuclei rep2 | 9868 | 63.3 % | 8 | 3 | 37.5 % | 1698 |
| 9. TotalSeq-A PBMC1 (healthy) | 14635 | 84.1 % | 1248 | 521 | 41.74 % | 754 |
| 10. TotalSeq-A PBMC2 (SARS-CoV-2) | 11372 | 83.6 % | 837 | 154 | 18.39 % | 721 |

**Sup. Table 4**. **Mislabelling ratios**. The mislabelling rate was calculated for each cell line by comparing the cells labelled as singlets by their specific hashtag (MULTISeqDemux) against their genotype (freemuxlet). The labelled cells belonging to a different genotype were considered mislabelled. The mislabelling % is presented as average values +/- SD across the 4 cell lines: MCF7, PC3, DU145, MDAMB231 (or 3 patients for PBMC samples).

| Hashing experiment | Mislabeling | SD |
|---|---|---|
| 1. TotalSeq-A cells | 0.1% | ± 0.03 |
| 2. TotalSeq-A cells rep2 | 0.18% | ± 0.15 |
| 3. TotalSeq-C cells | 0.11% | ± 0.13 |
| 4. LMO (MULTI-seq) cells | 0.89% | ± 1.1 |
| 5. LMO (custom) cells | 2.67% | ± 1.15 |
| 6. CMO nuclei | 0.12% | ± 0.05 |
| 7. TotalSeq-A nuclei | 4.07% | ± 2.51 |
| 8. TotalSeq-A nuclei rep2 | 3.81% | ± 4.24 |
| 9. TotalSeq-A PBMC1 (healthy) | 4.76% | ± 1.93 |
| 10. TotalSeq-A PBMC2 (SARS-CoV-2) | 1.53% | ± 1.06 |
